# Feature Importance Network reveals novel functional relationships between biological features in *Arabidopsis thaliana*

**DOI:** 10.1101/2022.05.15.492035

**Authors:** Jonathan Wei Xiong Ng, Swee Kwang Chua, Marek Mutwil

## Abstract

Understanding how the different cellular components are working together to form a living cell requires multidisciplinary approaches combining molecular and computational biology. Machine learning shows great potential in life sciences, as it has the ability to find novel relationships between biological features. Here, we constructed a dataset of 11,801 gene features for 31,522 *Arabidopsis thaliana* genes, and developed a machine learning workflow to identify linked features. The detected linked features are visualised as a Feature Important Network (FIN), which can be mined to reveal a variety of novel biological insights pertaining to gene function. We demonstrate how FIN can be used to generate novel insights into gene function. To make this network easily accessible to the scientific community, we present the FINder database, available at finder.plant.tools (http://finder.plant.tools/).

## Introduction

Recent advances in computational approaches and experimental workflows have made obtaining genome-wide biological and genomic data relatively easy and commonplace. The high-throughput data captures different biological features for DNA (e.g., sequence, methylation, chromatin accessibility, chromatic conformation) and RNA (e.g., sequence, abundance, structure, modification) for hundreds of plants^1^. However, the sheer volume of biological data presents a challenge for deriving biological meaning from it. As such, identifying how different biological features link to each other, and how they interact with genomic information remains a significant challenge^2, 3^ .

Until recently, the main approach to determine how biological features are linked would use complex statistical approaches that might be sensitive to the quality of the data^3^. Fortunately, machine learning has emerged as a popular technique in many biological contexts, to make predictions based on biological information fed into it. This is because some machine learning algorithms are efficient enough to handle massive data sets that exhibit high amounts of noise, dimensionality, and/or incompleteness, and make minimal assumptions about the data’s underlying probability distributions and generation methods^1^. Machine learning methods can usefully be segregated into two primary categories: supervised or unsupervised learning methods. Supervised methods are trained on labelled examples and then used to make predictions about unlabelled examples, whereas unsupervised methods find structure in a data set without using labels^4^.

Machine learning has been used to predict gene function in multiple contexts, such as predicting specialised metabolism genes^5^, and computational assignment of GO terms to genes^6–8^. For example, Moore et al.^5^ used around 10 000 features on a dataset of around 5 000 genes to predict specialised metabolism genes, achieving a true positive rate of 87% and a true negative rate of 71%. Kulmanov et al.^6^, Rifaioglu et al.^7^ and Littmann et al.^8^ used protein sequences as the sole data source for GO term prediction. A recent study by Cheng et al.^9^ used an evolutionarily-informed machine learning approach within and across species to predict genes affecting nitrogen utilisation in crops, and showed how their approach is also useful in mammalian systems.

Machine learning can be also used to infer gene regulatory networks. For example, a study showed that plant metabolism is transcriptionally coordinated via developmental and stress conditional processes^10^. Another approach, named EXPLICIT (Expression Prediction via Log-linear Combination of Transcription Factors), correctly predicted gene expression patterns from transcription factor information. EXPLICIT also enabled inference of transcription factor regulators for genes functioning in diverse plant pathways, including those involved in suberin, flavonoid, lateral root, xylem and the endoplasmic reticulum stress response^11^. Thus, with the ability to correctly predict gene function and regulation from high-dimensional data, machine learning has a great potential to transform biology.

However, while achieving accurate predictions is a key aim of machine learning, understanding which biological features contribute to making these predictions can reveal how the different biological features are linked. Fortunately, some machine learning approaches are interpretable, as they provide feature importance scores that quantify how important a feature is in predicting the target feature. When five categories of features, gene sequence, protein sequence, network topology, homology, and gene ontology-based features, were used to predict essential genes in *Caenorhabditis elegans*, the topology feature category provided the highest discriminatory power for essentiality prediction^12^. In using machine learning to predict *Arabidopsis thaliana* secondary metabolism genes, it was shown that multiple genetic features, such as tandem duplication, coexpression with paralogs, expression levels, conservation, and gene coexpression are predictive of secondary metabolism genes relative to general metabolism genes^5^.

Using reported regulatory pairs in *A. thaliana* along with gene expression and molecular information, Zaborowski et al.^13^ aimed to discern the molecular determinants of high expression correlation of transcription factors and their target genes. Specific molecular determinants such as transcription factor family assignment, stress-response process involvement, and young evolutionary age of target genes were found particularly indicative of high transcription factor target gene correlation.

The above examples showcase the power of machine learning in identifying information that explains the molecular wiring of plants. However, the above mentioned studies focused on specific aspects of gene function (essentiality, specialised metabolism and gene regulation), which precludes us from understanding how the different properties of genes are important for their function. To address this, we constructed an extensive dataset of 31 522 *A. thaliana* genes, drawn from 11 801 features from multiple biological and genomic categories. We then used a machine learning workflow on this dataset to test the predictability of all features, and observed that certain features are more predictable than others. Using feature importance values derived from our workflow, we constructed a Feature Importance Network (FINder), which can be used to study which of the 11801 features are putatively functionally related. To make our analyses publicly-available, we created an online database, finder.plant.tools (http://finder.plant.tools/). With FINder, we exemplify how potential novel biological relationships amongst features can be identified.

## Material and Methods

### Sequence information

Primary transcripts of *A. thaliana* coding sequences (CDS) and protein sequences are obtained from Phytozome (http://phytozome.jgi.doe.gov/pz/portal.html)^14^.

### Gene expression features

Gene expression levels as measured by transcript per million (TPM) values, and gene specificity measure (SPM) values were obtained from EVOREPRO^15^ (www.evorepro.plant.tools). Differential gene expression (DGE) features were obtained from RNA sequencing (RNA-seq) data from ArrayExpress^16^ (https://www.ebi.ac.uk/arrayexpress/), and processed with kallisto^17^ and sleuth^18^. Amplitude and time points of peak expression of diurnal genes, was downloaded from diurnal.plant.tools^19^ (https://diurnal.sbs.ntu.edu.sg/).

### Gene family, phylostrata and genomic information features

Gene families, defined as orthogroups, as well as phylostrata corresponding to each gene, were downloaded from EVOREPRO^15^. Smaller numbers (starting from 1) indicate older phylostrata and larger numbers (ending at 21) indicate younger phylostrata. Gene family size, single copy and tandemly duplicated genes were also identified.

### Protein domain and biochemical features

InterProScan 5.44-79.0^20^ on *A. thaliana* protein sequences was run, and the number of protein domains (Pfam), disordered regions (MobiDBLite) and transmembrane helices (TMHMM) were obtained. The total number of domains in each gene, protein length, isoelectric point (*pI*) and molecular weight of proteins were obtained, with the last two from the Isoelectric Point Calculator (IPC)^21^.

### Biological network features

Biological network features were made from Protein protein interaction (PPI) (BioGRID, https://thebiogrid.org/^22^), gene coexpression (EVOREPRO^15^), gene regulatory^13^, and functional gene networks (Aranet, http://www.inetbio.org/aranet/)^23^. For all networks, two network centrality measures, degree and betweenness centrality, were calculated. A markov cluster (MCL) algorithm^24, 25^ was used to cluster the PPI, gene regulatory and functional gene networks. The heuristic cluster chiselling algorithm (HCCA)^26^ was used to cluster the gene coexpression network through the calculation of highest reciprocal rank (HRR). HRR defines the mutual coexpression relationship between two genes of interest.

### Experimental GO terms as features

Gene annotations in the form of gene ontology (GO) terms were downloaded from The Arabidopsis Information Resource (TAIR, http://arabidopsis.org)^27^. Only gene annotations with experimental evidence codes EXP, IDA, IPI, IMP, IGI, and IEP were selected.

### Cis-regulatory element features

Cis-regulatory element names and families was downloaded from the Arabidopsis Gene Regulatory Information Server (AGRIS) database (https://agris-knowledgebase.org/)^28^. Their frequency per gene was calculated.

### Multi-omics (genomic and transcriptomic associated) features

Multi-omics features, in the form of GWAS and transcriptome-wide association studies (TWAS) were downloaded from the Arabidopsis thaliana multi-omics association (AtMAD) database^29^ (http://119.3.41.228/atmad/index.php). The number of times each gene was associated with each phenotype trait was counted.

### Evolutionary/conservation and epigenetic features

Homologous features were obtained from the EVOREPRO database^15^. Nucleotide diversity, methylation status of gene bodies, and sequence conservation features were obtained from a 2015 study by Lloyd, et al.^30^

### Protein post-translational modification (PTM) features

Protein PTM features were obtained from the Plant PTM Viewer^31^ (http://www.psb.ugent.be/PlantPTMViewer). For each gene, the number of each PTM together with the amino acid which it occurs in, was counted.

### Data preprocessing

All features were combined to create a dataset comprising 11 801 features and 31 552 genes for machine learning (Supplemental data 1). Missing categorical features were filled with 0. Missing continuous features were filled with their respective mean values, and continuous features were standardised (z-score normalisation). Care was taken to ensure only the training set was used to calculate these mean values and for the standardisation process, to prevent data leakage.

### Machine learning, time trial

To determine a suitable machine learning model which gives a good balance of performance and time taken to train, we tested logistic regression, random forest, balanced random forest, linear support vector machine (SVM) and adaboost.

To improve the model performance and estimate the time it takes to analyse all features, we set out to identify suitable hyperparameters. The range of hyperparameters tested for the time trial is given in Table 1.

**Table 1.** HPs tested for time trial.

| Model | Hyperparameter | Range of values |
| --- | --- | --- |
| Adaboost | n_estimators | 100, 120, 130, 150, 200 |
|  | learning_rate | 0.6, 0.625, 0.65, 0.675, 0.7, 0.725, 0.75, 0.775, 0.8 |
| Balanced random forest | max_features | sqrt, 0.1, 0.2, 0.3, 0.4, 0.5, 0.75 |
|  | n_estimators | 50, 100, 200, 500, 1000 |
|  | max_depth | 10, 20, 50, 70, 100, 125, 150, 200, 500, None |
| Logistic regression | C | 0.0001, 0.001, 0.01, 0.1, 1, 10, 100, 1000, 10000 |
| Linear SVM | C | 0.0001, 0.001, 0.01, 0.1, 1, 10, 100, 1000, 10000 |
| Random forest | ccp_alpha | 0, 0.1, 0.001, 0.001 |
|  | max_features | sqrt, 0.1, 0.2, 0.3, 0.4, 0.5, 0.75 |
|  | n_estimators | 50, 100, 200, 500 |
|  | max_depth | 20, 50, 100, 200, None |

We used a randomised search with 10 iterations within a nested 5-fold cross validation approach. In a k-fold nested cross validation approach, the inner k-fold is used for hyperparameter optimization during random search, while the outer k-fold fold is used to test the model, hence the equivalent of k test/validation sets would be used. Such an approach was used to train the model with hyperparameter optimization, so as to minimise data leakage (https://scikit-learn.org/stable/auto_examples/model_selection/plot_nested_cross_validation_iris.html). The F1 metric was used to score models due to unbalanced sample sizes between the positive and negative classes. Genes labelled with the specific feature used as the class label are in the positive class, while genes which do not have that feature, are in the negative class. Before model training, for the specific GO term used as the class label, all parent and child GO terms related to that class label were removed. Parent and child GO terms are identified using GOATOOLS^32^. Parent and child GO terms need to be removed to prevent data leakage during model training, as GO terms used as class labels are associated with their corresponding parent and child GO terms. This approach was also used for downstream machine learning applications, and modifications from this approach will be specified.

### Identification of optimal hyperparameters for random forest

To find the optimal hyperparameters for the random forest models, we chose 71 GO terms (Table S5), that contained between 5-1000 genes. The set of random forest hyperparameters tested is given in Table 1. The Out of Bag (OOB) F1 metric was used to score models. We used random search to estimate the optimal hyperparameters with 20 iterations for each GO term. We divided the OOB scores into three groups: high (OOB F1 >= 0.7), medium (0.5<= OOB F1 < 0.7) and low (OOB F1 < 0.5) scoring.

To identify hyperparameters that were frequently found in the high performing group, we used two approaches. In the first one, we selected the most frequently occurring hyperparameter value. In the second, we selected the most frequently occurring group of values. Hyperparameters chosen from these two methods were used to train random forest models for the 71 GO classes. Additionally, default hyperparameters, and hyperparameters optimised for each GO class were also used. The selected hyperparameter values were tested on both original and shuffled data obtained by randomly shuffling the dataset columns.

### Construction of models for 9535 features

To construct models for each feature, each feature would be selected as the prediction target and the remaining features would be used as the training features (the dataset). For GO terms, we removed child and parent features of a GO term that is used as a prediction target (label). GO terms with <10 genes were not used as targets. A total of 9535 features were used as prediction targets in our workflow. The OOB F1 score was the metric used for categorical features, while the OOB R^2^ score was the metric used for continuous features. We used the hyperparameters that resulted in best overall performance, as estimated in ‘Identification of optimal hyperparameters for random forest’: ccp_alpha = 0.001, max_features = 0.2, n_estimators = 50, and max_depth = 200.

### Feature importance network construction

Construction of the network utilised the concept of calculating mutual ranks between feature pairs. To remove poorly predictable features, only features which scored >= 0.4 with the OOB F1 (for categorical features) or R^2^ (for continuous features) metric were selected for mutual rank calculations, which left 1475 features. For these predicted features, all of their non-zero feature importance values were converted into ranks, with the feature with the largest feature importance value given a rank of one, and the feature with the smallest feature importance value given the largest rank. The mutual rank of each pair was then calculated by taking the geometric mean of both ranks, as described in ATTED-II^33^. The formula for calculating mutual ranks is given here:

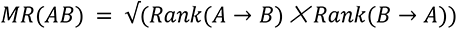

In the formula, MR stands for mutual rank, MR(AB) refers to the MR of features A and B, and Rank(A→B) refers to the feature importance rank which A has with respect to B. Rank(B→A) would refer to the inverse.

To find a MR cutoff, the top 10% of the MR values (53 080 MR values were obtained corresponding to 53 080 feature pairs) were used to build the network, comprising 1342 nodes and 5 308 edges. Edge weights were created by inverting a list of feature pairs, sorted by mutual ranks. An inversion is done since smaller mutual ranks indicate a stronger link between features, whereas a larger edge weight indicates the same idea. As such, the constructed network is a network of features, with putative biological links between them depicted by edges.

### Feature importance network analysis

The feature importance network was analysed to identify biologically relevant groups of features. Overall network metrics, which are betweenness centrality, clustering coefficient and degree, were calculated using Cytoscape 3.8.2^34^.

A permutation test was used to identify statistically significantly associated feature categories, which are defined as each row of the table given in Table 2. First, the number of edges between all possible feature category pairs was calculated. Next, the features in all feature category pairs were shuffled 10 000 times, and we compared the number of edges after shuffling, with the number of edges before shuffling (the original number of edges). The calculated empirical p-value indicates whether the original number of edges is significantly depleted as compared to random chance. P-values were corrected using the Benjamini-Hochberg correction^35^.

**Table 2.** Summary of features used. The first column describes the feature type. The second describes the feature name and parentheses indicate the number of features per name. The third column contains the feature description.

| Feature type | Feature name<br>(number of features) | Feature purpose |
| --- | --- | --- |
| Gene expression | SPM (9) | Expression specificity |
|  | TPM (6) | Gene expression levels |
|  | DGE (436) | Differential gene expression |
|  | Diurnal (13) | Diurnal gene expression, amplitude and time point |
| Gene family | Orthogroups (2) | Gene family size |
| Phylostrata | Phylostrata (1) | Phylostrata which genes belong to |
| Genomic information | Single copy genes (1) | Single copy genes in the same gene family |
|  | Tandemly duplicated genes (1) | Tandemly duplicated genes in the same gene family |
| Protein domain | MobiDBLite (1) | Prediction of disordered domains regions |
|  | Pfam (2761) | Collection of protein families |
|  | TMHMM (1) | Prediction of transmembrane helices |
|  | Number of domains (2) | Number of protein domains |
| Biochemical | Length of peptide (1) | Shows how long each peptide is |
|  | Molecular weight (1) | Molecular weight of peptide |
|  | Isoelectric point (pI) (1) | pI of peptide |
| PPI | Network centrality (2) | Degree and betweenness centrality |
|  | Network clusters (1295) | Cluster size and ID |
| Gene coexpression | Network centrality (2) | Degree and betweenness centrality |
|  | Network clusters (279) | Cluster size and ID |
| GO terms | GO terms (3645) | Experimentally determined gene annotations |
| cis-regulatory elements | cis-regulatory element names (82) | Gene regulation |
|  | cis-regulatory element families (15) | Gene regulation |
| Multi-omics | GWAS (33) | Genomic loci within genes, correlated with phenotype traits |
|  | TWAS (28) | Gene expression level, correlated with phenotype traits |
| Gene regulatory network | Network centrality (2) | Degree and betweenness centrality |
|  | Network clusters (55) | Cluster size and id |
|  | Properties (76) | Biological characteristics of transcription factors and their target genes |
| Aranet gene-interactions | Network centrality (2) | Degree and betweenness centrality |
|  | Network clusters (2957) | Cluster size and id |
| Evolution | Homologs (22) | Presence of *A. thaliana* homolog in 22 species |
|  | Nucleotide Diversity (1) | Nucleotide diversity calculated from *A. thaliana* accessions |
| Epigenetics | Gene body methylation (1) | Whether gene body is methylated |
| Conservation | Sequence conservation (3) | Protein sequence % identity to fungi, plants, metazoans |
|  | Percent identity to paralogs (1) | Maximum percent identity from BLAST to closest paralog |
|  | dN/dS values (4) | dN/dS substitution rates between *A. thaliana* |
|  |  | paralogs, and homologs from 3 plant species |
|  | Paralog dS (1) | dS with putative paralog |
| PTMs | Protein PTM (58) | Protein PTM frequency |

### Data analysis and availability

All data processing and analysis tasks, unless otherwise stated, used python 3.8.6 and its associated libraries for data science, such as pandas 1.1.4, numpy 1.19.4, scipy 1.6.1 and statsmodels 0.13.0. Networks were constructed and analysed using networkx 2.5^36^, and analysis and visualisation of the feature importance network also made use of Cytoscape 3.8.2^34^ and its associated apps. Machine learning was done by scikit-learn 0.23.2^37^ while the balanced random forest model was trained using imbalanced-learn 0.7.0^38^. Data visualisation was done using matplotlib 3.4.2^39^, seaborn 0.11.1^40^ and ptitprince 0.2.5^41^.

The machine learning dataset, together with scripts used, are available from a github repository (https://github.com/jonng1000/ml_plant). Raw data used to create this dataset is available upon request.

### Database development

The frontend is hosted on github and uses React.js and cytoscape.js^42^. The frontend code is available as a github repository (https://github.com/Sweekwang/golabel). The REST API backend uses python flask which retrieves data from Google Cloud Storage. The backend is hosted on Google App Engine.

## Results

### Assembly of 11 801 features for 31 552 Arabidopsis genes

We used 11 801 features for 31 522 *A. thaliana* genes, drawn from a wide variety of 37 categories (Table S1, Table 2). Gene expression features include SPM, TPM and differential gene expression (DGE) features. Gene SPM values (expression specificity, indicating whether a gene is e.g., specifically expressed in roots) were obtained from 9 organs, stem, female, male, leaf, flower, seeds, root, apical meristem, and root meristem. From TPM values from these organs, 6 summary statistics were calculated, which are mean, median, maximum, minimum, variance calculated by square of standard deviation and variance calculated by median absolute deviation divided by the median (MAD). DGE features were derived from 218 conditions, and each condition was used to create 2 features, which are the up and down regulated status for genes.

Genomic and evolutionary features include gene family (orthogroup), phylostrata, protein domains, gene regulation and homolog features. One type of gene family size was calculated by counting the number of *A. thaliana* genes in the gene family, whereas the second type was calculated by counting the number of genes from all species in the gene family. For each orthogroup, its last common ancestor was assigned as its phylostrata. One method of counting the number of protein domains counts the total number of domains in each gene, while the other counts the total number of unique domains. 76 features describing biological characteristics of transcription factors (TFs) and their target genes (TGs) were also obtained from the gene regulatory network. An *A. thaliana* gene is defined to have a homolog with a particular species if that species has a gene in the same orthogroup as that *A. thaliana* gene.

To conclude, we ended up with 9535 features as targets, as 2266 GO term features were removed as they had <10 genes per term. As such, 11 801 features were used in our machine learning workflow to predict 9535 targets.

### Finding the optimal machine learning model

To identify which machine learning method produces the most accurate predictions, we tested five methods (logistic regression, random forest, balanced random forest, linear support vector machines (SVM) and adaboost) and used the F1 score, which is the harmonic mean between precision and recall, to score the models. In addition, due to the resource intensiveness of the pipelines, we also noted the training time needed for finishing model training. For the time trial, 16 well annotated GO cellular component terms were used as labels (i.e., prediction targets, Table S2). Both logistic regression and linear SVM produced warnings as their algorithms could not converge with the specified number of iterations (a hyperparameter), thus their results were not reliable.

Therefore a second time trial was conducted, to determine the number of iterations needed for convergence and the time taken. 2 GO classes, GO:0016020 and GO:0005829, were used together with all five models, with 10 random search iterations and 5-fold nested cross validation. This would allow for a fair comparison of the time taken of all five models to be made, as a suitable number of iterations for logistic regression and linear SVM was used to ensure convergence. Using only 2 GO classes would help to ensure that all models were trained in a reasonable amount of time.

We observed that the random forest showed consistently high F1 scores (Figure 1A), and a reasonable amount of time to train (Figure 1B, Figure S1, Table S3). In addition, random forests allow one to use the out-of-bag (OOB) score to test the model, hence saving time by removing the need for a test/train split.

**Figure 1.**
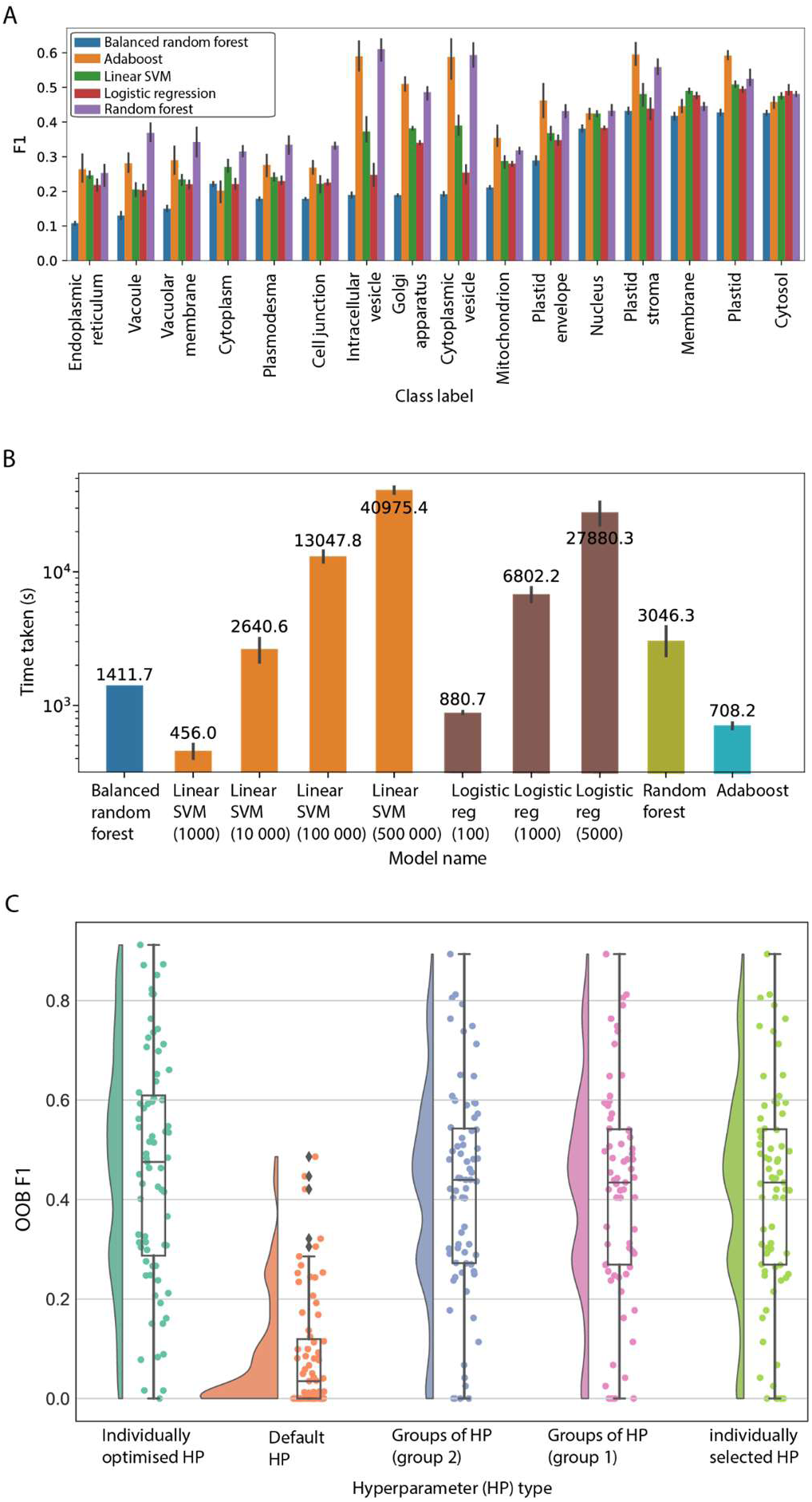
Evaluation of machine learning algorithms. A) F1 scores (y-axis) of 16 GO cellular location terms (x-axis). The algorithms are logistic regression (average F1 score 0.32), balanced random forest (0.26), adaboost (0.41), random forest (0.43) and linear SVM (0.35). The predictions were performed 5 times, and the error bars represent the 95% confidence interval. B) Time (y-axis) taken to train the different machine learning models to finish training. The predictions were performed 5 times on GO terms GO:0005829 (cytosol) and GO:0016020 (membrane). C) OOB F1 score (y-axis) for the 71 GO terms using different sets of hyperparameters. ‘Individually selected HP’ refers to random grid search to optimise hyperparameters for each GO term individually. ‘Default HP’ means that default hyperparameters are used. ‘Groups of HP (group 1)’ and (group ‘2)’, refers to the most frequent hyperparameter group observed after optimizing HPs for the 71 GO terms. ‘Most frequent individual hyperparameter’ refers to the most frequent individual hyperparameter chosen after optimizing for the 71 GO terms.

Machine learning models can be further tuned by adjusting their hyperparameters. To determine the suitable hyperparameters for the random forest model by taking into account model scores and training time, we tested the influence of four hyperparameters on the prediction performance of 71 GO labels (Table S5). These are cost-complexity pruning (ccp_alpha), maximum number of features for each split in the tree (max_features), number of trees in the forest (n_estimators) and maximum depth of the tree (max_depth). We investigated four methods to identify the best hyperparameters. The first method identified which individual hyperparameter values are most frequently found among the best-performing models of the 71 GO terms. The second method identified the most frequently occurring groups of hyperparameters. The third method used the default hyperparameters (which are ccp_alpha=0.0, max_features=auto, n_estimators=100, and max_depth=none). The fourth method used hyperparameters individually optimised for each GO term, which represents the most computationally-intensive approach to estimate the hyperparameters, as each model has to be optimised individually with cross-validation (Table S6 and Table S7).

The outcome of the analysis revealed that the individually optimised hyperparameters produced the best-performing models (Figure 1C, average F1 = 0.46). Conversely, the default hyperparameters performed the worst (Figure 1C, average F1 =0.09), showing that optimised hyperparameters can dramatically improve the performance of the models. Furthermore, two most frequent groups of parameters (group 1 and 2, average F1 scores of 0.41505 and 0.41536 respectively) and most frequent individual HP (F1 0.41504) performed comparably to the individually optimised HP (Figure 1C). These results show that ‘individually selected’ and ‘groups’ of hyperparameters perform very similarly to ‘individually optimised’ hyperparameters. Based on these results, an ‘individually selected’ hyperparameter values method was chosen to build models for each of the 9535 features. To verify that our random forest workflow was able to perform better than random chance, we used the found HPs on both original and randomly shuffled data. Results show that our workflow with the selected hyperparameters performs better than random chance (Figure S2) and hence was used for model training on all features.

### Calculating the predictive performance of biological features

To investigate which biological features can be predicted well by our machine learning model, for each of the 9535 features we built a random forest model. To score the performance of each model, we used the OOB F1 score for categorical features (0 and 1 represent poor and perfect performance, respectively), while for continuous features we used the OOB R^2^ score (< 0, 0 and 1 indicate predictions worse than always predicting the mean value of the target, poor performance [predict mean value of the target regardless of input] and perfect performance, respectively, Figure 2).

**Figure 2.**
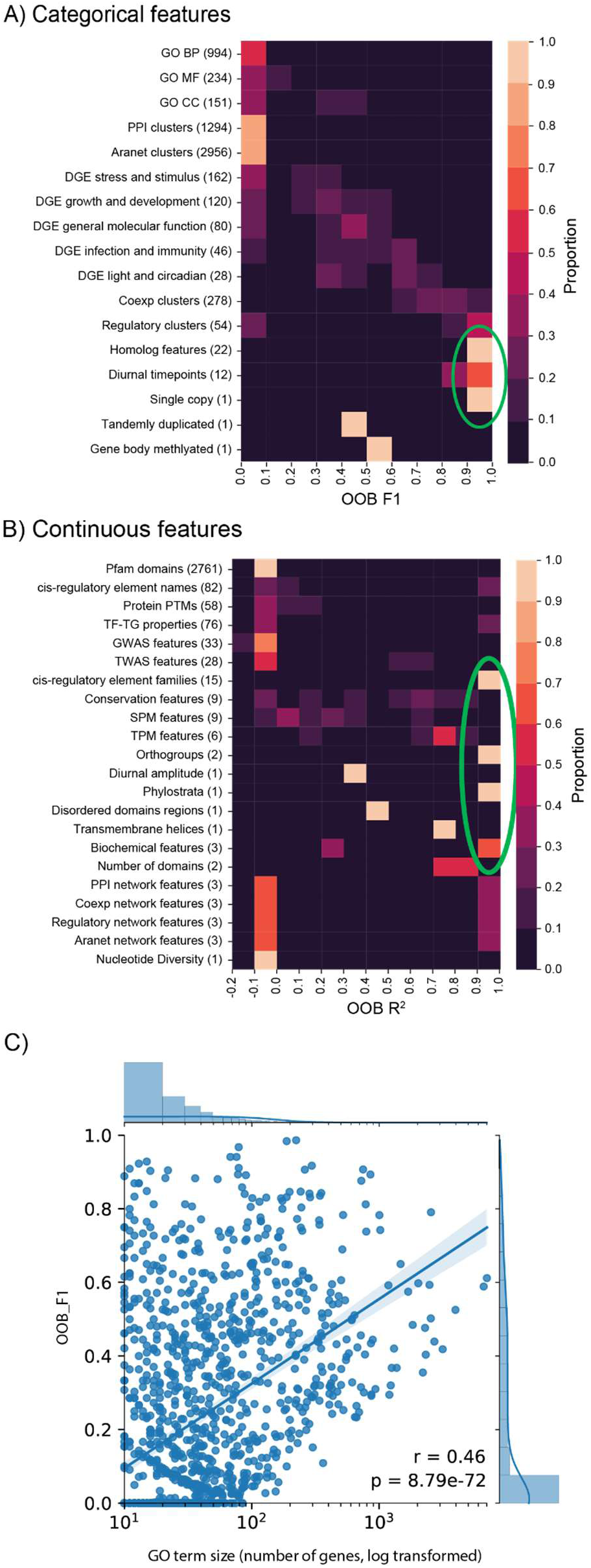
OOB F1 or R^2^ scores (x-axis) of the biological features. A) F1 scores (x-axis) for categorical features (y-axis). B) R^2^ continuous feature scores. C) F1 scores (y-axis) of GO terms vs. number of genes in each GO term (x-axis).

We set out to investigate how well the different types of features could be predicted. Features that could be predicted well (defined as lightly coloured squares to the right of the clustermaps and indicated by the green circles, in Figure 2A and 2B) comprise of homolog features, diurnal timepoints, single copy, cis-regulatory element families, orthogroups, phylostrata, and biochemical features. Conversely, features that could not be predicted well are all other features. While most GO terms and DGE are not accurately predicted, some of them have high scores (Figure 2A). To determine if the number of genes in each GO term influences model scores, a plot of scores against the number of genes was made (Figure 2C). This scatterplot showed a moderate but statistically significant relationship between the number of genes and OOB F1 score (Pearson’s r = 0.46, p-value = 8.79e-72). This indicates that increasing the number of genes in GO terms does positively influence model performance. Thus, the performance of machine learning models will be improved by the inclusion of more biological data for more genes.

### Construction of Feature Importance Network (FIN)

To investigate the biological relationships between features, we identified which features are mutually predictive of each other, and used this observation to infer biological relationships between them. To do this, we constructed a Feature Importance Network (FIN), where nodes represent machine learning features and edges represent features with putative biological relationships. To obtain the FIN, mutual ranks of feature pairs were calculated and the top 10% of them were used to construct the network. This is based on the assumption that these represent features that are mutually predictive of each other, which implies biological relationships between them. A total of 1 475 nodes (features) and 53 080 edges (mutual ranks) were present before applying the top 10% cutoff and after applying it, 1 342 nodes and 5 308 edges were selected to build the FIN.

To analyse the topology of the network, we first investigate the node degree of the FIN, and we observed a power-law distribution (FIgure 3A). This is consistent with a scale-free network topology observed in many biological networks such as metabolic^43^, RNA^44^, protein^45^ and gene coexpression^46^ networks. The distribution of the mutual ranks (Figure 3B) is similar to the HRR distribution of the gene coexpression data used in our study (Figure 3C). Given that gene coexpression networks are known to be scale-free^47^, this observed similarity can lend support to our inference of the scale-free nature of the FIN. Therefore, this network topology could imply that while many features are biologically linked to only a few others, a minority of features are biologically linked to many others.

**Figure 3.**
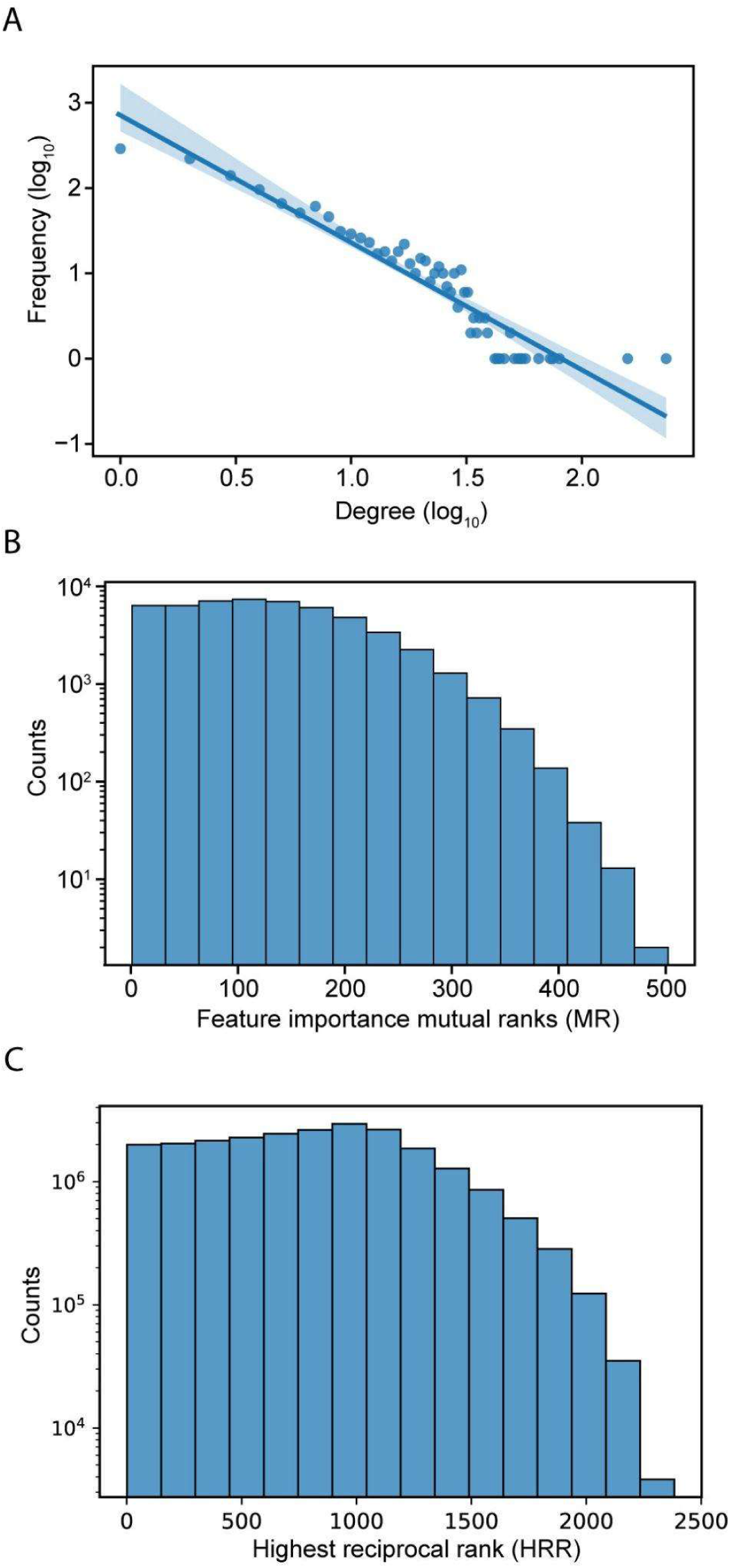
Characteristics of Feature Importance Network. A) Power-law distribution of node degrees. B) Distribution of mutual ranks (MR) of feature importance values. C) Distribution of highest reciprocal rank (HRR) of gene coexpression values in the coexpression network used in our study.

To visualise the FIN, we grouped the nodes (features) according to their corresponding categories (Table 2). While we observed many edges between the features of the same category, we also observed many edges between features from different categories (Figure 4). The complex web of interactions between the features show a complex relationship between the features of Arabidopsis genes.

**Figure 4.**
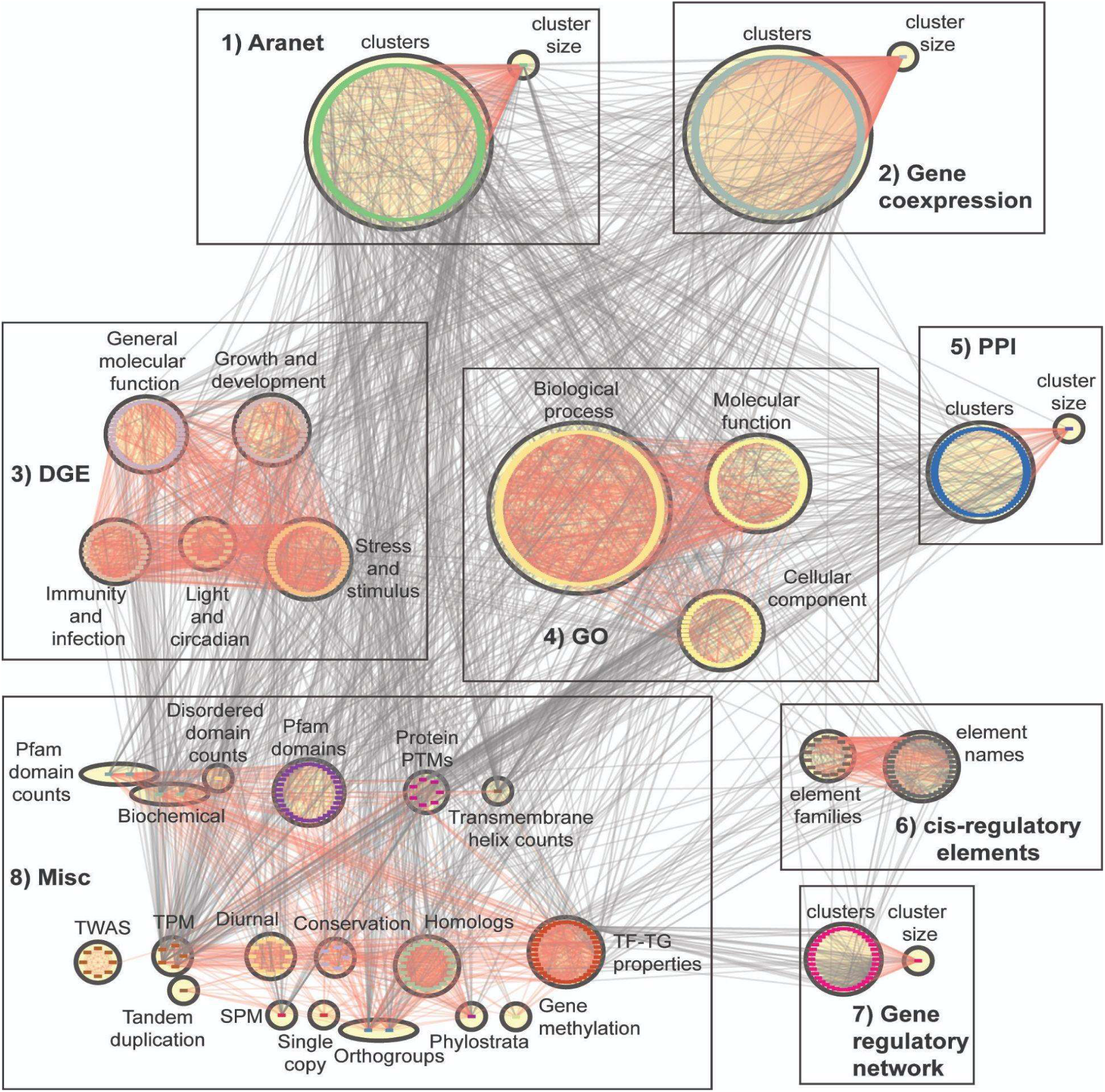
Feature importance network. Nodes represent features while edges connect features that have putative biological links. The features are divided into eight major groups (Table 2). Red edges show relationships within groups belonging to the same box. These groups are divided into eight boxes, and the first seven boxes represent clusters of feature categories. The boxes are 1 (Aranet features: cluster IDs and size), 2 (gene coexpression features: cluster IDs and size), 3 (DGE features: subdivided into five categories), 4 (GO terms: subdivided into three categories), 5 (PPI features: cluster IDs and size), 6 (cis-regulatory elements), 7 (gene regulatory features: cluster IDs and size), and 8 (a miscellaneous group which contains all other feature categories).

The node degree distribution of the feature categories in the FIN vary across categories (Figure 5). GO terms are observed to have low node degrees in general (top blue circle, Figure 5), while DGE features have a comparatively higher node degree distribution (bottom blue circle, Figure 5). This implies that GO terms tend to be less functionally linked to other features, while DGE features exhibit more functional links. Orthogroup and phylostrata (top yellow circle), transmembrane helices, biochemical features (length and molecular weight of peptide) and number protein domains (middle yellow circle), and network features (bottom yellow circle), are feature categories which exhibit high node degree distributions (Figure 5). This implies that they share functional links with many other features. The network features with a high node degree distribution are the cluster size feature of their respective biological networks, which are connected to many cluster ID features (Box 1,2,5 and 7 in Figure 4).

**Figure 5.**
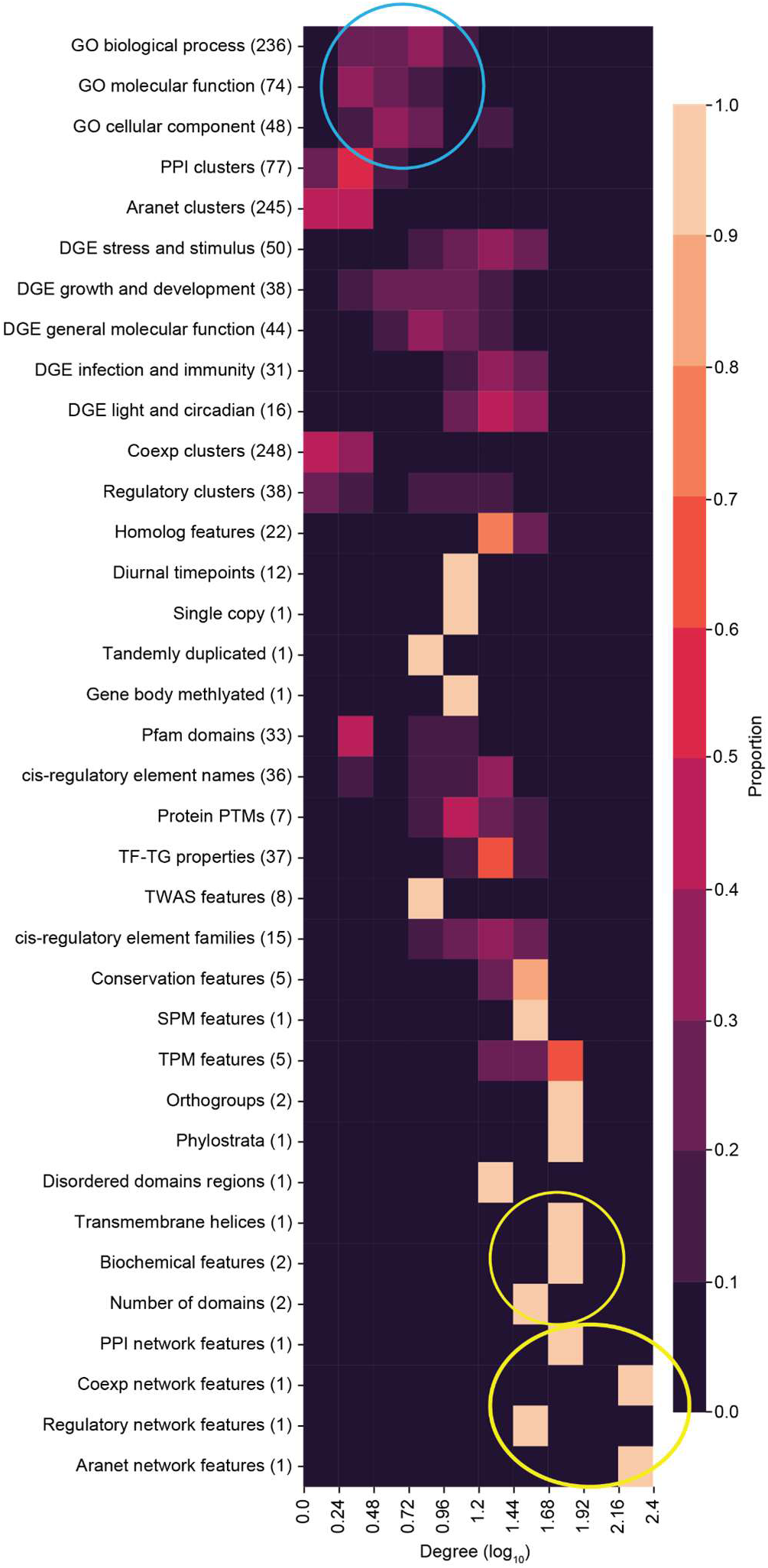
Degree distribution of feature categories. Top blue circle shows GO terms while the bottom blue circle shows DGE features. Top yellow circle shows orthogroups and phylostrata, middle yellow circle shows transmembrane helices, biochemical features (length and molecular weight of peptide) and number protein domains, and bottom yellow circle shows network features (cluster size) from the PPI, coexpression, regulatory and Aranet networks.

### Identification of functionally-related feature categories

To identify which feature categories are significantly linked to each other, we determined which categories have more edges between them than expected by chance. This was achieved by performing a permutation test on the number of edges linking categories from the FIN together, and comparing the permuted number with the original number of edges (Figure 6).

**Figure 6.**
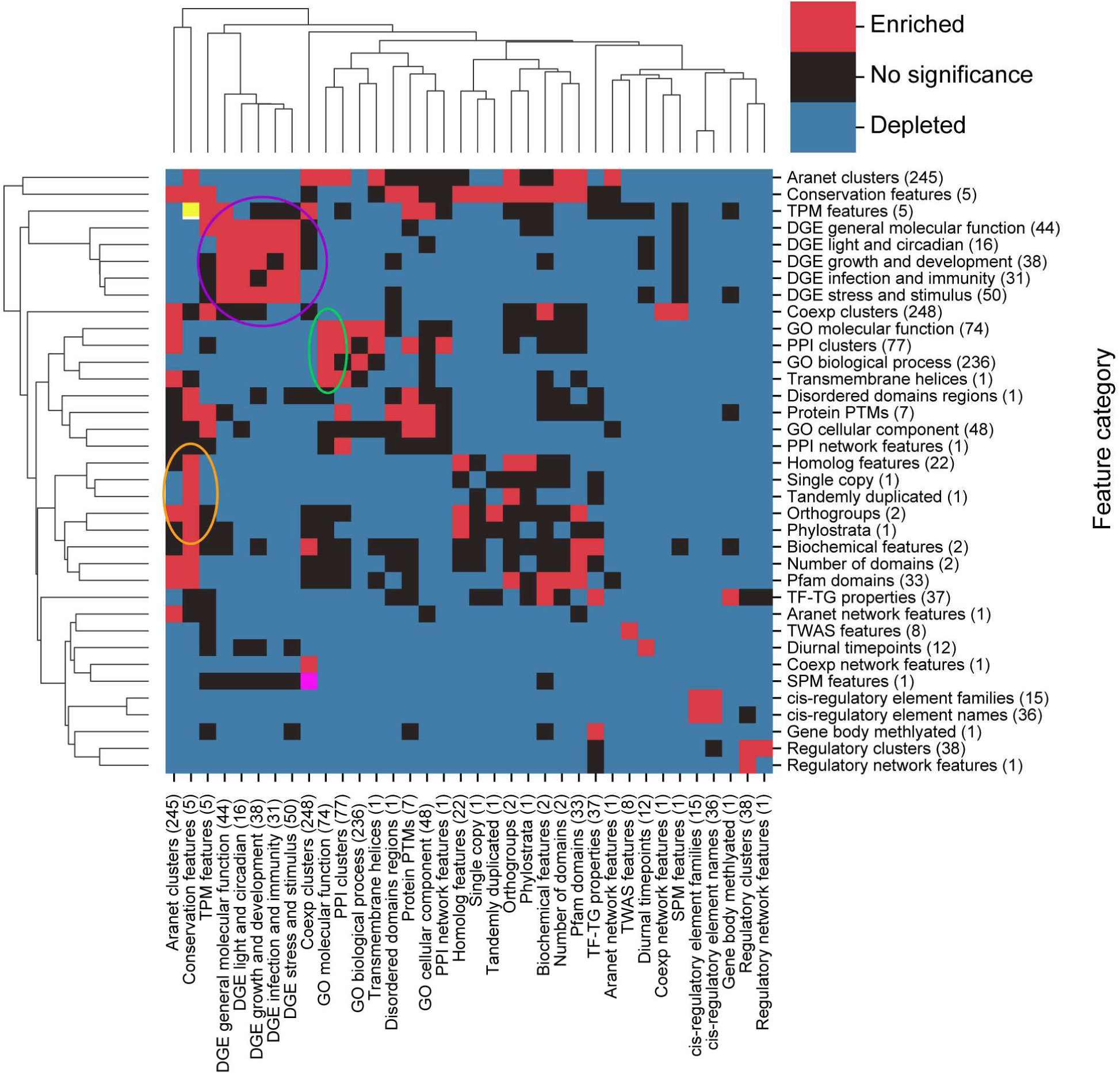
Identification of significantly associated feature types. The clustermap shows whether there are significantly (BH-adjusted p-values < 0.05) more (red squares) or significantly less (blue square) between feature categories that are expected by chance. Not statistically significant associations are indicated by black boxes. Circles indicate clusters of feature categories which are associated with each other. Yellow (top left) and pink (bottom left) squares are red squares which have been coloured differently to highlight them.

Most pairs of feature categories have a significantly smaller number of connections between them than expected by chance (Figure 6, blue squares), indicating that they are less likely to be functionally related. The exceptions are five DGE groups (purple circle), GO molecular function, PPI clusters, GO biological process and transmembrane helices (green circle), conservation features (conservation of gene sequences), homologs, single copy, tandemly duplicated, orthogroups, and phylostrata (orange circle), that represents clusters of significantly connected feature categories (Figure 6). Conservation and TPM features (yellow square, top left in Figure 6) and coexpression clusters and SPM features (pink square, bottom left in Figure 6) are pairs of features which are also seen to be linked.

The number of biological relationships within feature categories also tend to be higher than expected, as depicted by the many red squares along the clustermap diagonal. Taken together, features tend to share functional links within the same category, compared to across categories.

### Construction of finder.plant.tools - online database to browse FIN

To provide a user-friendly interface for scientists to explore the FIN and identify biologically associated features, we created an online database (http://finder.plant.tools/), finder.plant.tools, that can be queried by feature names. The database can display the local FIN neighbourhoods of features. To represent positively or negatively correlated features, and the strength of the correlations, we use red edges to indicate positive association, and blue edges indicate negative association. Grey edges indicate associations between neighbours which do not involve the target feature. The edge width indicates the strength of association (based on mutual rank), where a thicker width indicates a stronger association.

To demonstrate the ability of the FIN database to capture biologically-relevant information, we set out to investigate several features with expected associations. First, we observed that mean gene expression (tpm_mean, brown rounded rectangle) is positively associated with maximum (tpm_max, brown rounded rectangle) and median (tpm_median, brown rounded rectangle) gene expression (Figure 7A). This indicates that genes with higher mean expression tend to have higher median and maximum expression. Conversely, we observed a negative association between mean gene expression, and protein molecular weight (pep_mw, green rounded rectangle) and length (pep_aal, green rounded rectangle). This indicates genes with lower mean expression, tend to code for proteins with a lower molecular weight and have a shorter length. Furthermore, we observed that genes belonging to co-expression clusters 24 and 254 (cid_cluster_id_24 and 254, blue rounded rectangles), genes belonging to the ribonuceloprotein complex (GO_CC_ribonucleoprotein complex, beige rounded rectangle) and genes involved in glutathione binding (GO_MF_glutathione binding, yellow rounded rectangle), tend to have lower mean expression.

**Figure 7.**
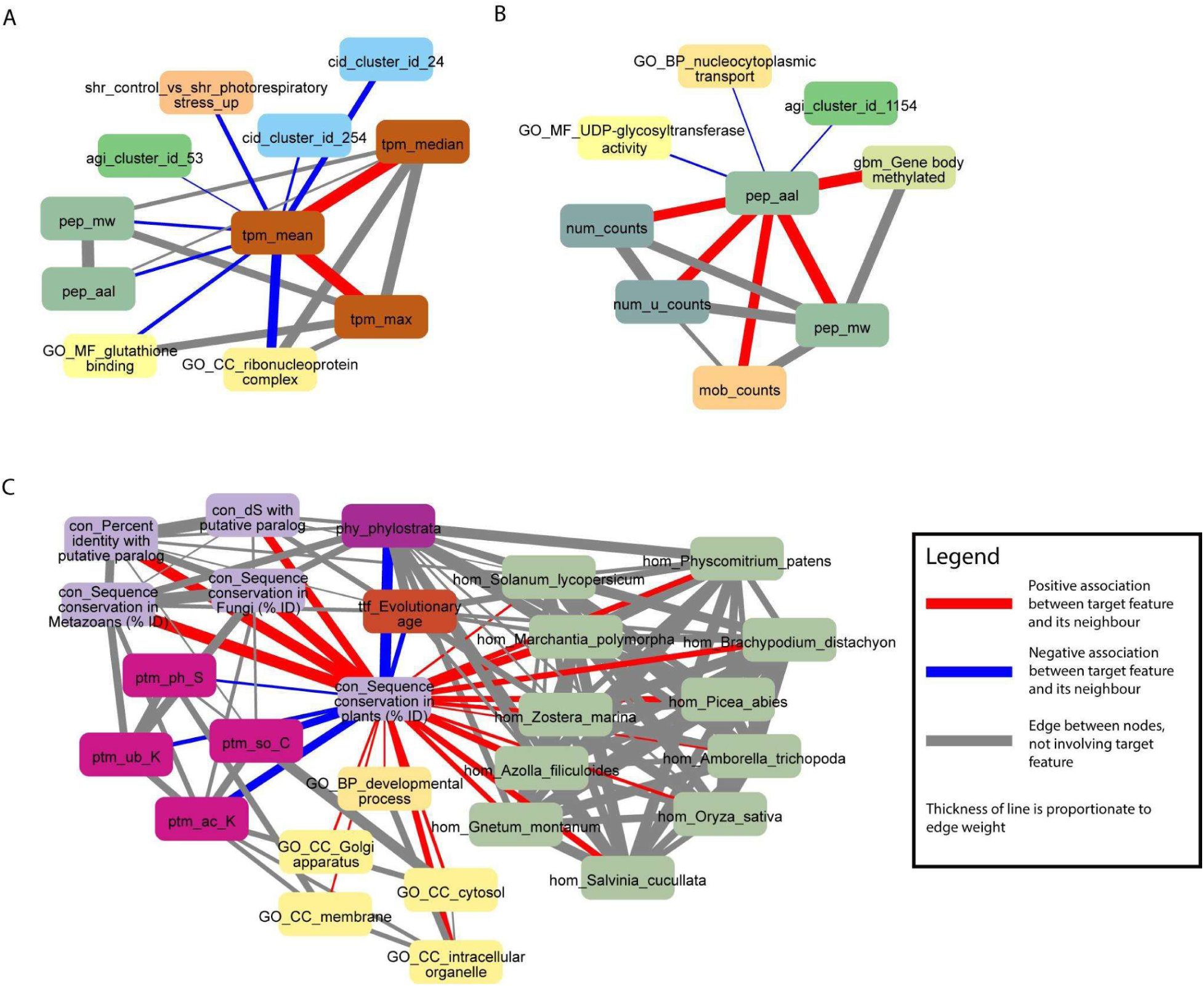
Examples capturing protein sizes and conservation. Selected nodes and edges from the local neighbourhood of specific features in the database are shown. A) Mean gene expression (tpm_mean) is positively associated with maximum (tpm_max) and median (tpm_median) gene expression. B) Protein length (pep_aal) is positively associated with the number of protein domains (num_counts), number of unique protein domains (num_u_counts), number of disordered protein domains (mob_counts) and protein molecular weight (pep_mw). C) Sequence conservation in plants (con_Sequence_conservation_in_plants_(%_ID)) is positively associated with sequence conservation in paralogs, fungi and metazoans (nodes starting with “con_”), fundamental GO terms (nodes starting with “GO_”) and homology with multiple plant species (nodes starting with “hom”). Sequence conservation in plants is negatively associated with the evolutionary age of target genes of transcription factors (ttf_Evolutionary_age), phylostrata (phy_phylostrata) and protein PTMs (nodes starting with “ptm_”).

In the second example we observed that protein length is positively associated with the number of protein domains (num_counts, greenish beige rounded rectangle), number of unique protein domains (num_u_counts, greenish beige rounded rectangle), number of disordered domains (mob_counts, beige rounded rectangle), and protein molecular weight (Figure 7B). Not surprisingly, proteins with a longer length tend to have a higher number of protein domains, more disordered domains and a higher molecular weight. However, we also observed a link between methylation of the gene body and protein length, suggesting that longer proteins tend to be more regulated on the epigenetic level.

Finally, in the third example, we saw that sequence conservation in plants (con_sequence conservation in plants (% ID), light purple rounded rectangle) is positively associated with sequence conservation in paralogs (con_dS with putative paralog and con_percent identity with putative paralog, light purple rounded rectangle), fungi (con_sequence conservation in fungi (% ID), light purple rounded rectangle) and metazoans (con_sequence conservation in metazoans (% ID), light purple rounded rectangle), and homology with multiple plant species (green rounded rectangles on the right) (Figure 7C). Conversely, sequence conservation in plants is negatively associated with the evolutionary age of genes captured by phylostrata (phy_phylostrata, dark purple rounded rectangles) (Figure 7C). Phylostrata feature ranges from 1 (gene families found in Archaeplastida) to 21 (gene families only found in *A. thaliana*), and the negative association between sequence conservation and phylostrata can be explained by the fact that genes with higher conservation are older, and thus have a lower phylostrata number. Interestingly, sequence conservation in plants is negatively associated with protein post-translational modifications (PTM, ptm_ph_S: serine phosphorylation, ptm_so_C: cysteine S-sulfenylation, ptm_ub_K: lysine ubiquitination, ptm_ac_K: lysine acetylation, purple rounded rectangles), indicating that younger proteins tend to be most post-translationally modified. Taken together, these examples reveal a mixture of expected and novel insights, indicating that the FIN can be used to gain new knowledge about the molecular wiring of Arabidopsis.

We set out to investigate more complex examples. First, we observed that post- translational modification of lysine acetylation is positively associated with multiple GO cellular locations (e.g. GO_CC_thylakoid, GO_CC_chloroplast stroma, and GO_CC_chloroplast, beige rounded rectangles), especially those related to the chloroplast (Figure 8A), implying that lysine acetylation takes place in chloroplast, or is important for importing proteins into the chloroplast. Furthermore, we observed associations to Aranet cluster 70 (agi_cluster_id_70, green rounded rectangle) and PPI cluster 1 (pid_cluster_id_1, blue rounded rectangle), suggesting that proteins with this post-translational modification tend to physically interact.

**Figure 8.**
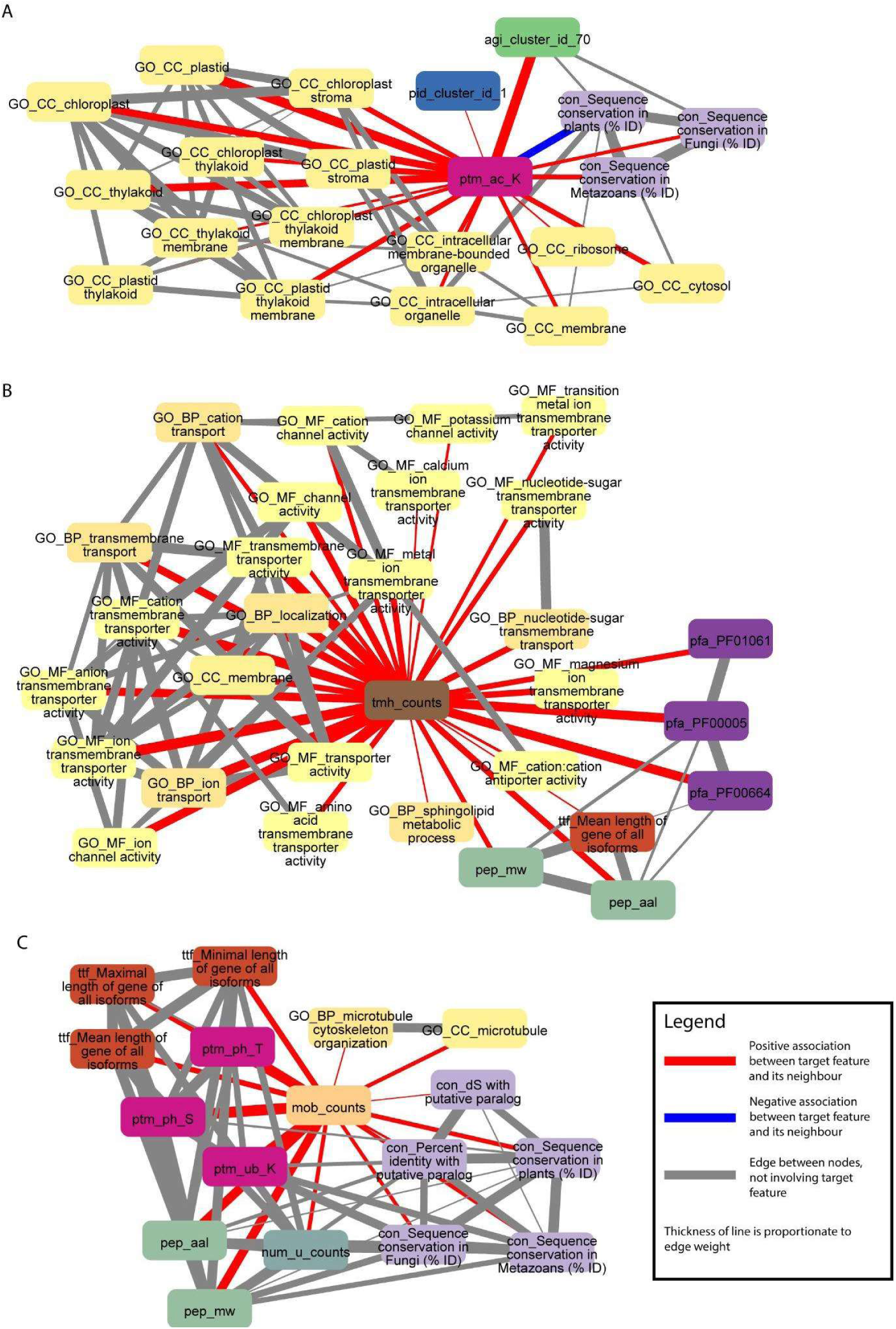
Examples capturing posttranslational modifications and the number of transmembrane and disordered domains. Selected nodes and edges from the local neighbourhood of specific features in the database are shown. A) Protein PTM - lysine acetylation (ptm_ac_K) is positively associated with GO cellular components terms (nodes starting with “GO_CC_”), especially those related to the chloroplast. B) Number of transmembrane helices in a protein sequence (tmh_counts) is positively associated with GO transmembrane transporter and channel terms (nodes starting with “GO_”), and sphingolipid metabolic process (GO_BP_sphingolipid metabolic process). C) Number of disordered domains in a protein sequence (mob_counts) is positively associated with sequence conservation in paralogs, plants, fungi and metazoans (nodes starting with “con_”), GO microtubule terms (nodes starting with “GO_”), gene length (nodes starting with “ttf_”), protein PTMs (nodes starting with “ptm_”), protein length (pep_aal), number of unique protein domains (num_u_counts), and protein molecular weight (pep_mw).

In the second example, we observed that the number of transmembrane helices in a protein sequence (tmh_counts, brown rounded rectangle) is positively associated with multiple GO terms associated with channels (e.g. GO_BP_transmembrane transport, GO_MF_cation channel activity and GO_MF_potassium channel activity, beige/yellow rounded rectangles), not surprisingly indicating that channels tend to have more transmembrane helices (Figure 8A). In addition, we also observed association to three Pfam domains (pfa_PF01061: ABC-2 type transporter, pfa_PF00005: ABC transporter, pfa_PF00664: ABC transporter transmembrane region, purple rounded rectangles), which is in line with proteins with transporter domains having more transmembrane helices. Furthermore, there is a strong correlation between the number of transmembrane helices, molecular weight and protein length, which can be explained by proteins having more domains (transmembrane helices, disordered domains, others), being longer. Interestingly, we also observed an association to the sphingolipid metabolic process (GO_BP_sphingolipid metabolic process, beige rounded rectangles), suggesting that this metabolic process involves proteins with transmembrane helices (Figure 8B).

In the third example, we observed that the number of disordered domains in a protein sequence is positively associated with sequence conservation in paralogs, plants, fungi and metazoans, suggesting that proteins with many disordered domains are of an ancient origin (Figure 8C). In line with the above examples in Figure 8B, the number of disordered domains is correlated with protein size (pep_aal: protein length, pep_mw: protein molecular weight, num_u_counts: number of unique protein domains). Interestingly, the number of disordered domains is positively correlated to several posttranslational modifications (ptm_ph_T: threonine phosphorylation, ptm_ph_S: serine phosphorylation, ptm_ub_K: lysine ubiquitination, purple rounded rectangles), and the number of unique protein domains a protein has (num_u_counts). The latter suggests that multi-domain proteins tend to contain disorganised domains. Finally, we observed that of all biological processes captured by GO, microtubules tend to be the only one’s positively associated with the number of disordered domains.

## Discussion

Understanding biological relationships between molecular characteristics is critical to understanding how life works, and machine learning has great potential to contribute to achieving such an objective. Here, we analysed 31 522 *A. thaliana* genes using 11 801 features. These features are drawn from a wide variety of categories such as genomic, transcriptomic, evolutionary, biochemical, and protein and gene interactions. We applied a machine learning workflow using random forests, and constructed a FIN that shows features with putative biological relationships.

We observed that it is important to optimise hyperparameters for increased machine learning performance as the models gained on average a 4.6-fold increase in performance after optimisation. Interestingly, rather than optimising parameters for each of the 9535 models individually, which is computationally costly, hyperparameters optimised for a subset of 71 models resulted in similar performance. Since individual optimization takes a significantly longer time for training compared to using fixed hyperparameters, we chose the most frequently occurring, individually selected hyperparameter values for model training.

While the performance of most GO term and DGE feature models were not high, a minority had high scores (Figure 2A). We observed a significant positive correlation between the number of genes in each GO term and OOB F1 score (Figure 2C). This implies that poor machine learning performance could be due to the small number of genes in many of the features. A study by Rifaioglu et al.^7^ also observed a strong correlation between GO term size and performance. While studies predicting GO terms have reported higher scores^6–8^, these studies focused on GO terms with a larger number of genes through inclusion of computationally annotated GO terms as prediction targets^8^.

We observed that the FIN shows a power-law distribution (Figure 3), indicating that it is a scale-free network, which is typical for many biological networks, such as protein, metabolic and coexpression networks^48^. We observed that many features are connected to each other, highlighting the complex web of biological interactions involved in the molecular wiring of Arabidopsis (Figure 4). DGE features have a comparatively higher number of functional links to other features than GO terms (Figure 5), and many of them tend to be fellow DGE features (Figure 4). This suggests that different stimuli can activate similar differential gene expression programs, which is the basis for online tools such as AtCAST^49^ (http://atpbsmd.yokohama-cu.ac.jp/cgi/atcast/home.cgi). For example, the diverse stress factors that plants face often activate similar cell signalling pathways and cellular responses, such as the production of stress proteins and upregulation of the antioxidant machinery^50^. Genes belonging to different orthogroups and phylostrata have been shown to be associated with organ-specific gene expression, gene functions such as cell cycle organisation and phytohormone action, and diverse abiotic stress responses^15, 51^.

To provide access to the FIN, we constructed an online database, FINder, available at http://finder.plant.tools/. The examples generated by FINder showed expected biological relationships between features. For example, mean gene expression is positively associated with maximum and median gene expression (Figure 7A), which is expected as mean and median are measures of central tendency, hence both of them are expected to be positively associated. Furthermore, protein length is positively associated with the number of protein molecular weight, protein domains, and disordered regions (Figure 7B), which is expected as longer proteins can contain a greater number of domains and disordered regions. Furthermore, sequence conservation in plants is positively associated with sequence conservation in paralogs, fungi and metazoans, fundamental GO terms and homology with multiple plant species (Figure 7C). This is an expected finding as the greater the degree of sequence conservation in plants, the more likely that the gene sequence is conserved throughout evolution. Furthermore, highly conserved genes are more likely to play key essential and fundamental functions, such as in developmental processes^51, 52^.

Interestingly, sequence conservation in plants is negatively associated with serine phosphorylation, cysteine S-sulfenylation, lysine ubiquitination, and lysine acetylation (Figure 7C). This indicates that younger proteins tend to be more posttranslationally modified than older proteins, which is supported by studies suggesting that the range of PTMs has increased throughout evolution^53, 54^.

Further analysis of posttranslational modifications showed that lysine acetylation is positively associated with multiple GO cellular locations, especially those related to the chloroplast (Figure 8A). Lysines are found in the subcellular localization signal domains of proteins, and their acetylation can regulate protein subcellular localization^55, 56^. Lysine acetylation may be an important posttranslational modification in the chloroplast, as four Calvin cycle enzymes are acetylated^57^. Studies in strawberry^58^, soybean^59^, rice^60^, tea leaves^61^ and wheat^62^, indicate that a large proportion of lysine-acetylated proteins are predicted to be localised to the chloroplast. Therefore, lysine acetylation associations identified by finder.plant.tools support these studies, and suggest that it plays a key role in chloroplast function.

We also observed associations between PTMs (threonine and serine phosphorylation, lysine ubiquitination) with the number of disordered domains (Figure 8C), which is in line with disordered regions in proteins being posttranslationally modified^63, 64^. The proportion of PTM sites was recently shown to be higher in the intrinsically disordered protein domains than the structured domains,^65^ where phosphorylation of serine and threonine, acetylation, and methylation were over-represented in disordered regions of seven species (animals, plants, fungi). Interestingly, lysine ubiquitination is another PTM which we observed (Figure 8C), that to the best of our knowledge has not been documented in the literature as being tied to disordered regions.

## Conclusion

To conclude, we created a dataset of 11 801 features with 31 552 *A. thaliana* genes, and used machine learning to propose functional links between the features. Feature importance values from our approach were used to create a Feature Importance Network (FIN), which revealed a variety of potentially significant biological relationships between different types of features. An online database, finder.plant.tools (http://finder.plant.tools/), was created to provide a user-friendly way of accessing the FIN.

## Supplementary data

**Supplemental data 1: Machine learning dataset, tarball file**

**Table S1. Complete feature table.**

**Table S2. 16 GO terms, along with their description, used for testing the scores of 5 ML models**

**Table S3. Summarising results of the first time trial experiment.**

**Table S4. Table showing the number of classes chosen for hyperparameter optimisation, note that a few of them have < 5 chosen when there is < 5 which did not meet the criteria.**

**Table S5. Table showing 71 GO terms, along with their description, used for hyperparameter optimization of random forest, to determine which is the best hyperparameter set for ML workflow development.**

**Table S6. Frequency of individual HPs during HP optimisation across different score categories, excel sheet.**

**Table S7. Frequency of groups HPs during HP optimisation across different score categories, excel sheet.**

## Supporting information

Supplemental talbe

---

**Figure S1.**
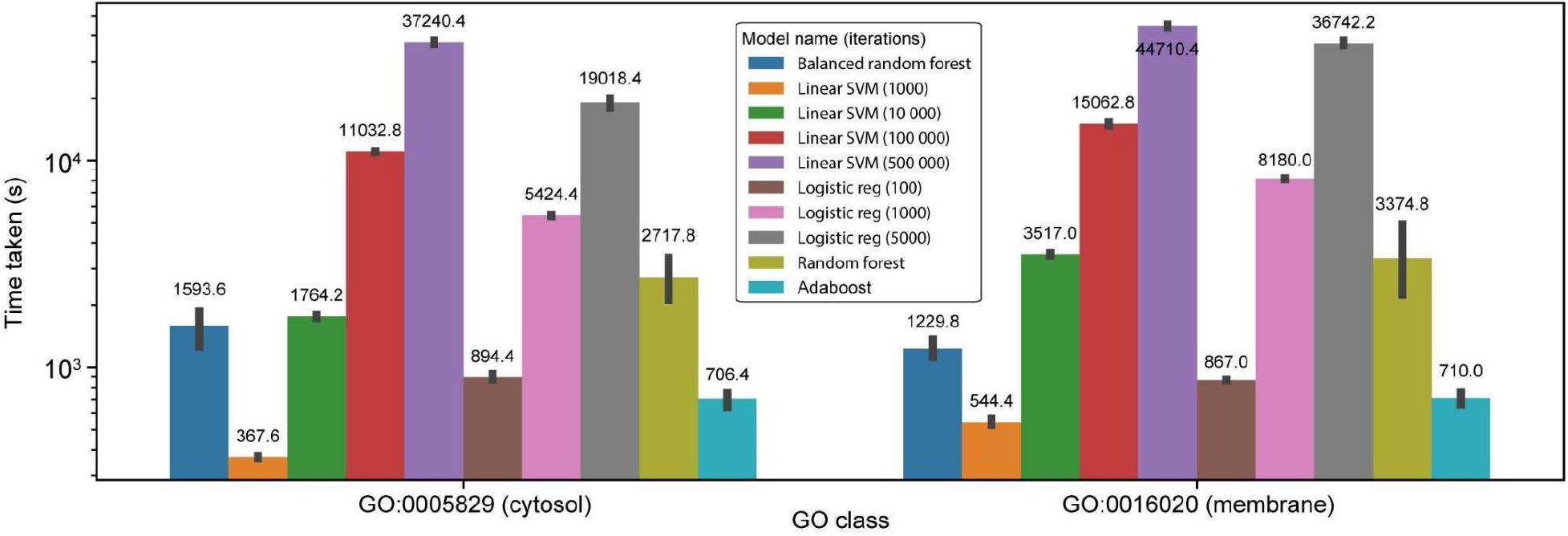
Shows time taken for each individual GO class. Time trial repeat - running all machine learning models on 2 GO classes, 5 repeats per model, to allow for all models to be compared on a common basis.

**Figure S2.**
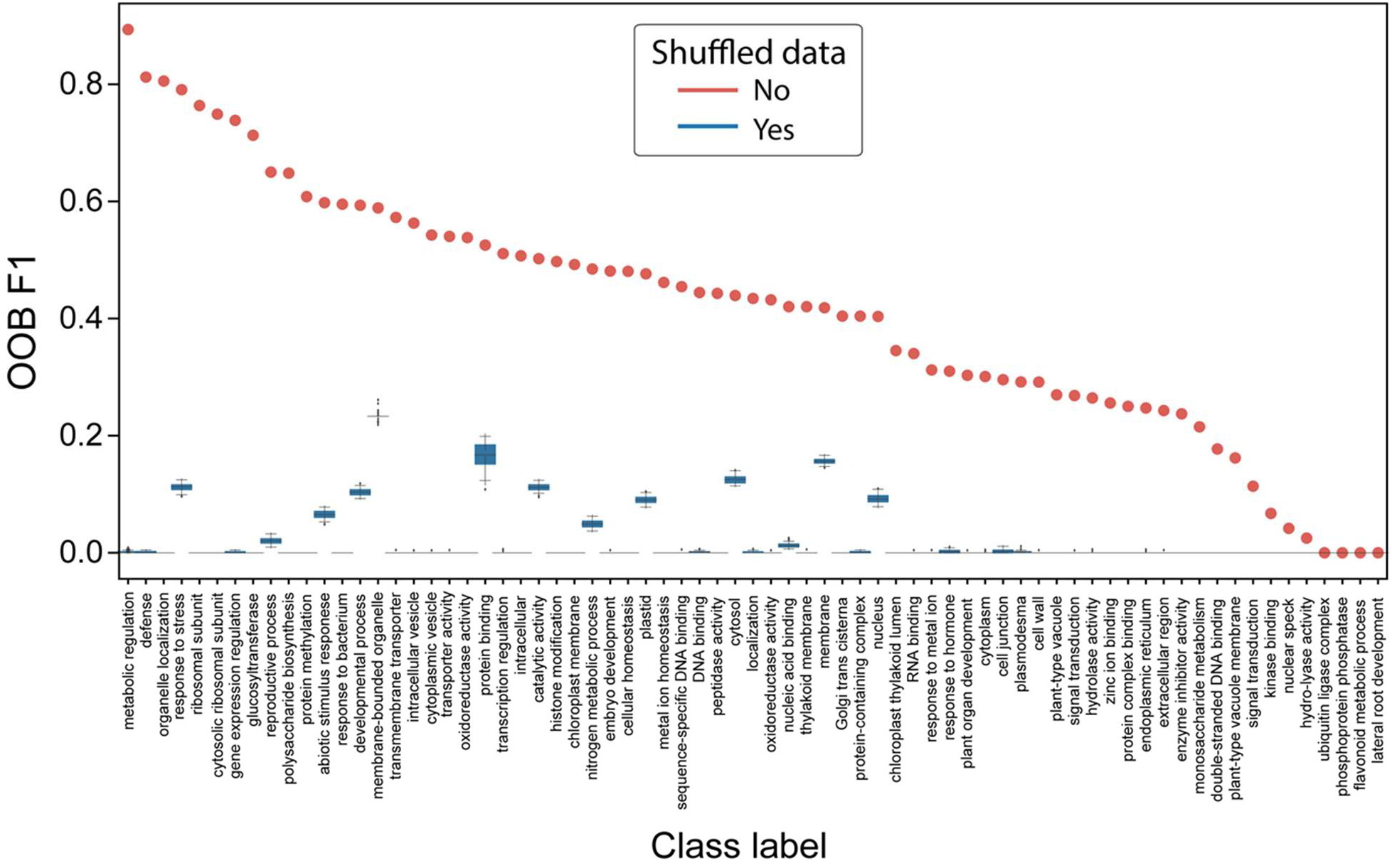
Random forest OOB scores when trained on shuffled data (blue) or actual data (orange). Some GO term descriptions have been summarised to save space, full descriptions are given in Table S9.

**Figure S3.**
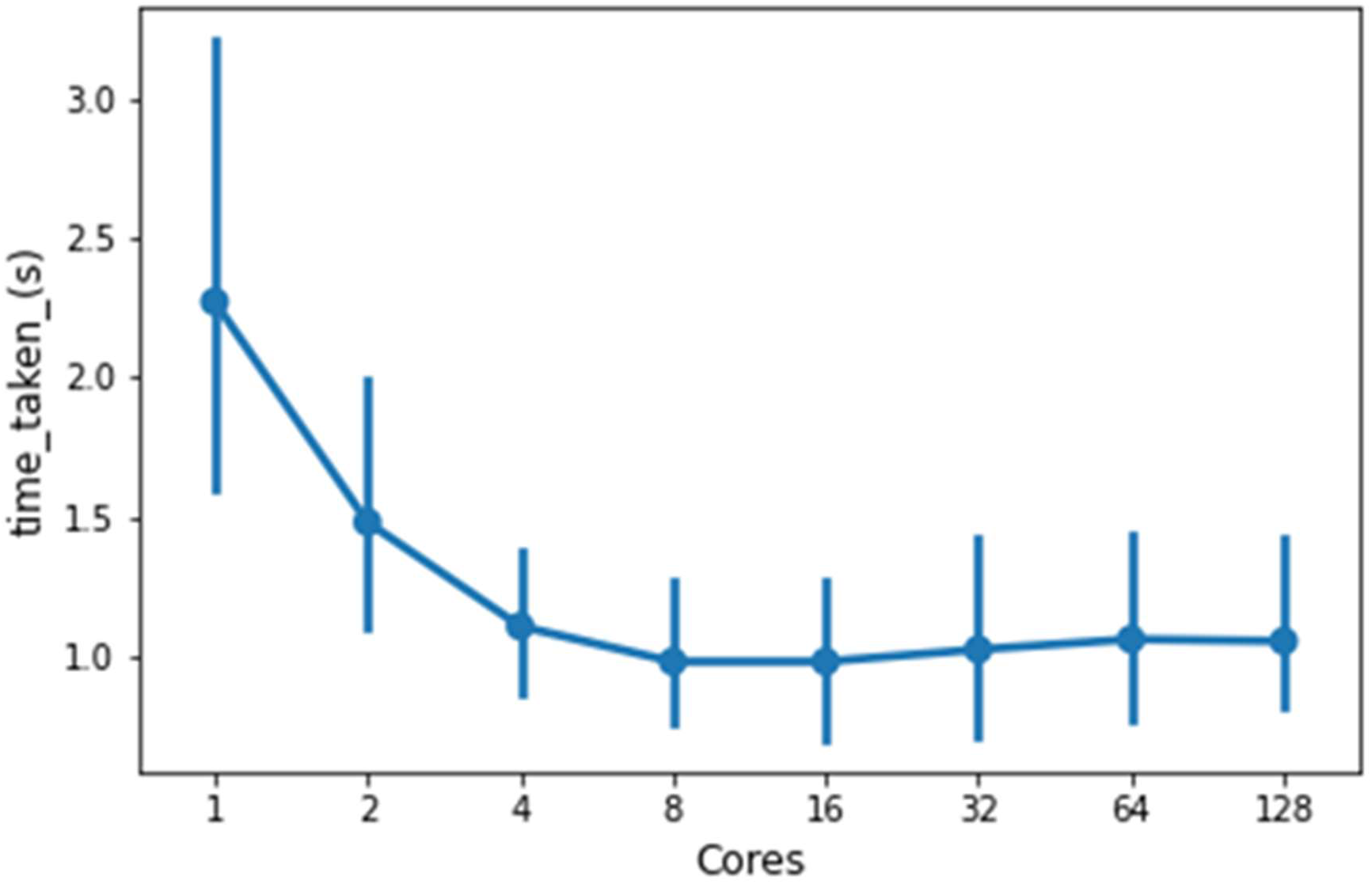
Line plot showing time taken to train random forest models on 5 GO classes, GO:0043229 (intracellular organelle), GO:0005774 (vacuolar membrane), GO:0048827 (phyllome development), GO:0030001 (metal ion transport) and GO:0016836 (hydro-lyase activity).

**Figure S4.**
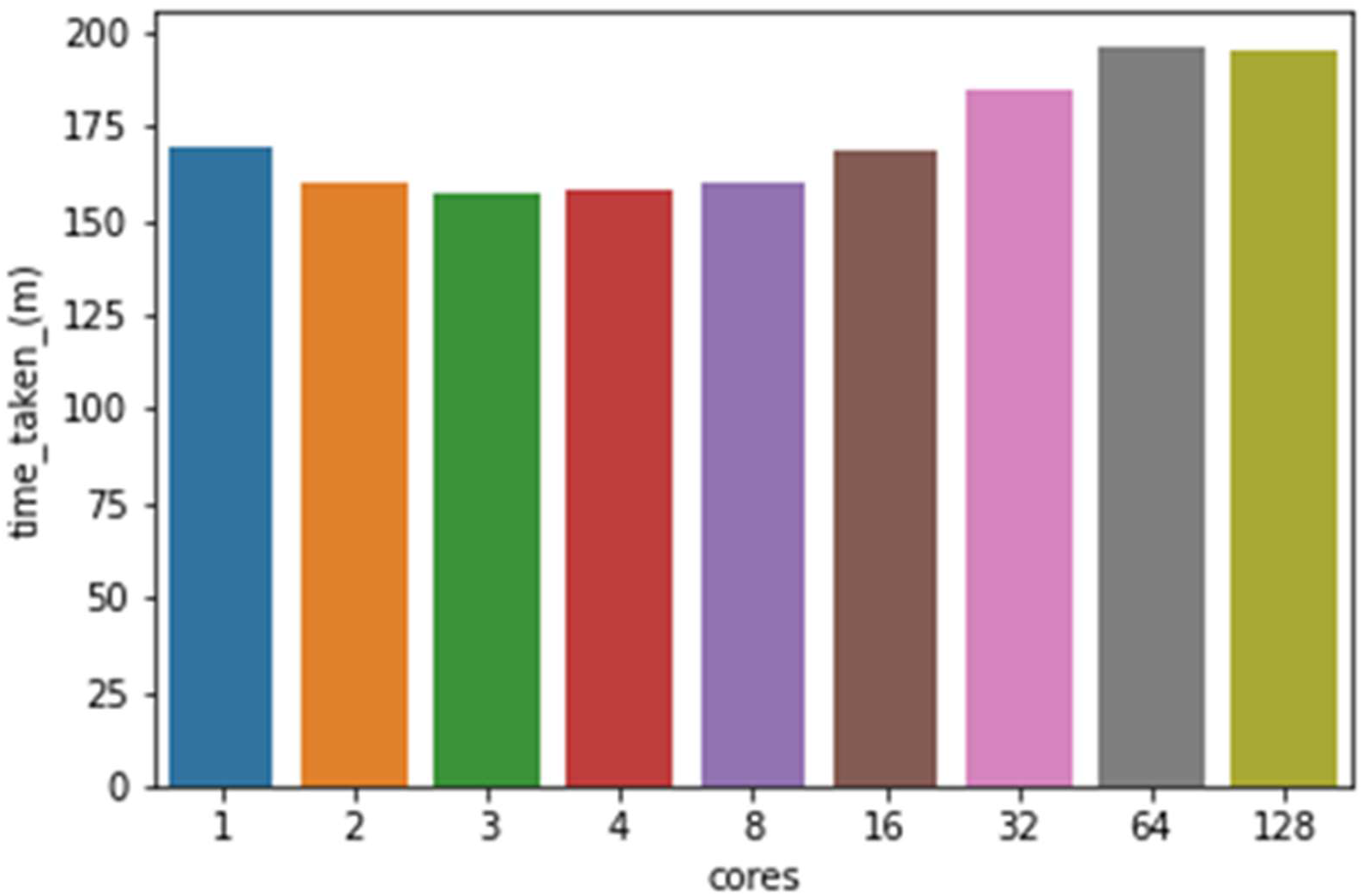
Bar graph showing time taken to train random forest models on all GO classes.

## Notes

http://finder.plant.tools

### Competing Interest Statement

The authors have declared no competing interest.

