## Supplemental Methods for "Feature Importance Network reveals novel functional relationships between biological features in *Arabidopsis thaliana*"

### Supplementary methods

#### Gene expression features

Gene expression levels as measured by transcript per million (TPM) values, and gene specificity measure (SPM) values were obtained from EVOREPRO, a published online database^15^ ([www.evorepro.plant.tools](http://www.evorepro.plant.tools)) which contains genomic and transcriptomic data derived from 13 members of the plant kingdom. Gene expression TPM and gene SPM values were obtained from 9 organs, stem, female, male, leaf, flower, seeds, root, apical meristem, and root meristem. From these TPM values, 6 summary statistics were calculated. These are mean, median, maximum, minimum, variance calculated by square of standard deviation and variance calculated by median absolute deviation divided by the median (MAD). Variance by MAD was calculated as this could be a useful summary statistic to use for gene expression values^30^. SPM values indicate the organ specificity of gene expression values, which ranges from 0 (not expressed in an organ) to 1 (expressed only in the organ). 9 SPM values are obtained. TPM and SPM values are continuous features.

Differential gene expression (DGE) features were obtained from collecting RNA sequencing (RNA-seq) datasets from ArrayExpress^16^ (<https://www.ebi.ac.uk/arrayexpress/>). All suitable RNA-seq datasets in ArrayExpress from the earliest dataset to July 2020 were downloaded. Suitable datasets are defined as those with RNA-seq data for at least one control and test condition. They also needed to have at least triplicate RNA-seq samples per condition. These datasets were downloaded from the European Nucleotide Archive (ENA) as fastq files. The RNA-seq data was then mapped against the CDS of *A. thaliana* and quantified using kallisto^17^. Kallisto quantifications were then analysed with the R package sleuth^18^, which identifies genes which have been differentially expressed. DGE analysis was done by comparing gene expression between test and control conditions. 218 conditions were obtained from ArrayExpress, and each condition was used to create 2 features, which are the up and down regulated status for genes. Hence, a total of 436 DGE features were obtained, which are categorical features, as a gene would be given a value of 1 if it was up/down regulated and 0 if it was not.

Diurnal gene expression data was downloaded from Conekt^19^ (<https://conekt.sbs.ntu.edu.sg/>), which is a published database which visualizes diurnal plant gene expression. The JTK_CYCLE algorithm^66^ was used to identify genes with significantly (adjusted P-value < 0.05) rhythmic expression patterns from the gene expression matrices. Amplitude of diurnally expressed genes was used as a continuous feature. 12 peak timepoints, each with a 2 h interval in a 24 h day, were used as categorical features. These refers to diurnal genes with maximum expression in a 24 h day. A value of 1 means a gene’s expression peaked at a specific timepoint, and a value of 0 means it did not.

#### Gene family and phylostrata features

Gene families, defined in this work as orthologous gene groups (orthogroups), as well as phylostrata corresponding to each gene, were downloaded from EVOREPRO^15^. Orthogroups were obtained using Orthofinder v.2.4.0^67^. The age of orthogroups (phylostrata) was estimated using phylostratigraphy^68^. For each orthogroup, its last common ancestor was assigned as its phylostrata, and given a numerical value, where smaller numbers (starting from 1) indicate older phylostrata and larger numbers (ending at 21) indicate younger phylostrata.

Gene family size was calculated using two different methods, to produce two gene family size features. One type of gene family size was calculated by counting the number of *A. thaliana* genes in the gene family, whereas the second type was calculated by counting the number of genes from all species in the gene family. Both gene family size and phylostrata are continuous features.

#### Genomic information features

Single copy genes refer to whether or not in one orthogroup, only one *A. thaliana* gene from its species is present. If this is true, that gene is said to be present as a single copy and given a value of 1, and if it is false, that gene is not a single copy and given a value of 0. Orthogroup and genomic information were used to identify tandemly duplicated genes. Genomic information was obtained from the Arabidopsis_thaliana.TAIR10.44.gff3 file downloaded from EnsemblPlants release 44^69^. For each gene in each gene family, if there are genes in the same family that are adjacent (defined as no other genes in between) to it in the chromosome, that gene would be considered as tandemly duplicated, and the gene is assigned a value of 1. If that is not the case, the gene would be considered to be not tandemly duplicated, and the gene is assigned a value of 0. Both single copy and tandemly duplicated features are categorical features.

#### Protein domain features

Protein domain features were created by running InterProScan 5.44-79.0^20^ on *A. thaliana* protein sequences. The Pfam, MobiDBLite and TMHMM database hits were extracted from InterProScan results. The number of each Pfam protein domain was counted for each gene, and domains which only appear in one gene were removed. Hence, 2761 continuous Pfam domain features were created. MobiDBLite is a database which predicts disordered domains regions in proteins, thus it is a continuous feature showing the number of disordered domains regions in *A. thaliana* genes. TMHMM is a database which predicts transmembrane helices in proteins, thus it is a continuous feature showing the number of transmembrane helices in *A. thaliana* genes.

The number of protein domains were counted via two methods, hence two continuous features were created. One method counts the total number of domains in each gene, while the other counts the total number of unique domains.

#### Biochemical features

Protein length was calculated based on *A. thaliana* protein sequences, and this was used as a continuous feature. Isoelectric point (*pI*) and molecular weight of *A. thaliana* protein sequences were calculated using the Isoelectric Point Calculator (IPC)^21^. *pI* was obtained from the IPC_Protein method in IPC as the IPC paper showed it to be the most accurate method. Both *pI* and molecular weight are continuous features.

#### Biological network features

Protein protein interaction (PPI) network information was downloaded from the BioGRID database 4.0.189 (https://thebiogrid.org/)^22^. Two network centrality measures, degree and betweenness centrality were calculated. These form two continuous features. A markov cluster (MCL) algorithm^24,25^ was used to cluster the PPI network into PPI clusters, and each cluster was given an identification (ID) number. Cluster IDs were one hot encoded to indicate whether each gene was found in each cluster ID. Cluster IDs with only one gene were removed. Hence, a total of 1294 cluster IDs as categorical features were obtained. Cluster size, a continuous feature, was also calculated.

Gene coexpression network information was derived from gene expression TPM values from the EVOREPRO database^15^. From these gene expression values, a highest reciprocal rank (HRR) based Pearson correlation approach was used to create the network^26^. A HRR < 100 cutoff was used as this seemed to us, a reasonable compromise between richness of information and false positives in identifying coexpressed genes. Two network centrality measures, degree and betweenness centrality were calculated. These form two continuous features. The network was clustered using the heuristic cluster chiselling algorithm (HCCA)^26^ and cluster IDs were one hot encoded to indicate whether each gene was found in each cluster ID. Cluster IDs with <5 genes were removed, hence a total of 278 categorical features were obtained. Cluster size, a continuous feature, was also calculated.

Gene regulatory network features were obtained from a paper identifying correlated expression of transcription factors (TF) and their target genes (TG)^13^. Two network centrality measures, degree and betweenness centrality were calculated. These form two continuous features. A markov cluster (MCL) algorithm^24,25^ was used to cluster the network into clusters, and cluster IDs were one hot encoded to indicate whether each gene was found in each cluster ID. Cluster IDs with only one gene were removed. Hence, a total of 54 cluster IDs as categorical features were obtained. Cluster size, a continuous feature, was also calculated. 76 continuous features, describing biological characteristics of TFs and TGs were also obtained.

Functional gene network features were downloaded from the Aranet database (http://www.inetbio.org/aranet/)^23^, from the AraNet.txt file. Two network centrality measures, degree and betweenness centrality were calculated. These form two continuous features. A markov cluster (MCL) algorithm^24,25^ was used to cluster the network into clusters, and cluster IDs were one hot encoded to indicate whether each gene was found in each cluster ID. Cluster IDs with only one gene were removed. Hence, a total of 2956 cluster IDs as categorical features were obtained. Cluster size, a continuous feature, was also calculated.

#### Experimental GO terms as features

Gene annotations in the form of gene ontology (GO) terms were downloaded from the ATH_GO_GOSLIM.txt file located in The Arabidopsis Information Resource TAIR (http://arabidopsis.org)^27^. Only gene annotations with experimental evidence codes EXP, IDA, IPI, IMP, IGI, and IEP were selected. These experimental GO terms were one hot encoded to indicate whether each gene was annotated with each term. GO terms with only one gene were removed. Hence, 3645 GO terms as categorical features were created. GO terms from the go.obo file were also downloaded from the GO database^70,71^, which were used in downstream development of our machine learning workflow.

#### Cis-regulatory element features

Cis-regulatory element information was downloaded from the Arabidopsis Gene Regulatory Information Server (AGRIS) database (https://agris-knowledgebase.org/)^28^. This information was divided into cis-regulatory element names and cis-regulatory element families, and the number of these names and families appearing in each gene was counted. Thus 82 continuous cis-regulatory element name features, and 15 continuous cis-regulatory element family features were created.

#### Multi-omics (genomic and transcriptomic associated) features

Multi-omics features, in the form of GWAS and transcriptome-wide association studies (TWAS) were downloaded from the Arabidopsis thaliana multi-omics association (AtMAD) database^29^ (<http://119.3.41.228/atmad/index.php>). The GWAS information from AtMAD shows the correlation between phenotypic traits with genomic loci within genes, and 33 phenotypic traits were available. Hence, the number of times each gene was associated with each phenotype trait was counted, which produced 33 continuous features. The TWAS information from AtMAD shows the correlation between phenotypic traits with gene expression level, and 28 phenotypic traits were available. Hence, the number of times each gene was associated with each phenotype trait was counted, which produced 28 continuous features.

#### Evolutionary features

Homologous features were obtained from the EVOREPRO database^15^. An *A. thaliana* gene is defined to have a homolog with a particular species if that species has a gene in the same orthogroup as that *A. thaliana* gene. EVOREPRO has orthogroups from 23 plant and algae species, including *A. thaliana*^15^. Homologous features are one hot encoded to indicate whether each gene has a homolog with each species, hence 22 categorical features are created.

Nucleotide diversity, a continuous feature, was obtained from a 2015 study by Lloyd, et al.^30^ In that study, nucleotide diversity was calculated from 80 *A. thaliana* accessions.

#### Epigenetic features

The methylation status of gene bodies was used as a categorical feature, where 0 means the gene body was not methylated and 1 means it was. This feature was obtained from a 2015 study by Lloyd, et al.^30^

#### Conservation features

Conservation features were also obtained from a 2015 study by Lloyd, et al.^30^ Sequence conservation, defined as protein sequence percentage identity for each *A. thaliana* gene to fungi, plants, metazoans was used as 3 continuous features. Percent identity to paralogs, defined as the maximum percent identity from BLAST to closest paralog, for each *A. thaliana* gene, was used as a continuous feature. Paralogs were identified in *A. thaliana*, rice, and *Saccharomyces cerevisiae*. Nonsynonymous (dN)/synonymous (dS) substitution rates between A. thaliana paralogs, and homologs from 5 plant species were also obtained from the study by Lloyd, et al.^30^ However, there was extremely limited information on dN/dS substitution rates from homologs from 2 plant species, hence these 2 species were dropped. In total, 4 continuous features in the form of dN/dS substitution rates were obtained. Paralog dS, defined as dS with putative paralog for each *A. thaliana* gene, a continuous feature, was obtained as a feature.

#### Protein post-translational modification (PTM) features

Protein PTM features were obtained from the Plant PTM Viewer ^31^(http://www.psb.ugent.be/PlantPTMViewer). For each gene, the number of each PTM together with the amino acid which it occurs in, was counted. This resulted in 58 continuous features being made.

#### Identification of optimal hyperparameters for random forest

Suitable hyperparameters were determined by training random forest models on a set of GO terms. Since there is a great variety in the number of genes annotated with each GO term, the set of GO terms used for hyperparameter optimization was drawn from GO terms across the three GO domains (biological process, molecular function, cellular component) with between 5-1000 genes. Table S4 shows how many GO terms were chosen per GO term gene size and GO domain. The aim was to choose five GO terms per gene size for each GO domain. However, less than five GO terms would be picked if less than five were available given these requirements. A total of 71 GO terms were selected for hyperparameter optimization and their names are given in Table S5. The set of random forest hyperparameters tested is shown in Table 1. OOB F1 was used to score models, and is the metric used for downstream machine learning work. Optimisation of hyperparameters was done using 20 iterations of random search.

After hyperparameter optimization was done, hyperparameters contributing to high model scores were identified (Table S6 and Table S7). This was done by dividing all trained models into three groups, high scoring groups (OOB F1 >= 0.7), medium scoring groups (0.5<= OOB F1 < 0.7) and low scoring groups (OOB F1 < 0.5). A relatively high threshold of 0.7 was used to identify high scoring groups as we wanted to be strict on the threshold to identify good hyperparameters. From the high scoring groups, different methods were used to identify the best performing hyperparameters.

#### Optimising number of cores

Our workflow was run on a custom built Ubuntu workstation, with 64 AMD EPYC™ 7551 cores running at 2 GHz (128 processors are available), and 750 GB RAM. Due to the fact that sci-kit learn allows some models, like random forest, to be trained using multiple cores, the optimal number of cores for our workflow was identified. This was done to ensure that our workflow could be run efficiently, given that an increase in the number of cores may not lead to a proportionate decrease in time taken. Initially 5 GO terms, GO:0043229 (intracellular organelle), GO:0005774 (vacuolar membrane), GO:0048827 (phyllome development), GO:0030001 (metal ion transport) and GO:0016836 (hydro-lyase activity), were chosen to measure the time taken for model training on them. 1, 2, 4, 8, 16, 32, 64, and 128 processes per GO term, were used in this experiment (Figure S3). A repeat of this experiment was conducted, with the same range of processes per GO term, but this time, all GO terms were used for model training (Figure S4). 4 cores were selected to be used in our downstream workflow to train all models.

#### Feature importance network construction

Random forest models are somewhat interpretable as they produce feature importance values for each feature used in predicting the target. These values indicate the degree to which a certain feature influenced the prediction, with a higher value indicating that the feature was more important in prediction.

Construction of the network utilised the concept of calculating mutual ranks between feature pairs. Only features which scored >= 0.4 with the OOB F1/R^2^ metric were selected for mutual rank calculations as we took this set of features to be those which could be predicted to some extent by machine learning. This score threshold was lower than the >= 0.7 used to identify high scoring groups for hyperparameter selection, as our results (Figure S2) showed that randomly shuffled data produced scores of 0 - 0.2. Hence we decided that a score of >= 0.4 indicated model performance which was significantly better than random, and was thus suitable for our work. For these predicted features, all of their respective feature importance values were converted into ranks, with the feature with the biggest feature importance value given a rank of one, and the feature with the smallest feature importance value given the largest rank. For each predicted target, because they will have features with feature importance values of 0, these features would be removed and not converted into ranks. Thus ranks are only assigned to features if they have non-zero feature importance values for their respective prediction targets. A matrix of ranks would hence be created, where the columns represent predicted features, and the rows represent all features in the dataset, and values in the matrix would be ranks.

This matrix was converted into a column of all possible feature pairs, where each pair would be associated with two ranks. For example, the pair of features A and B would be associated with the rank of B with respect to A, and the rank of A with respect to B. The mutual rank of each pair was then calculated by taking the geometric mean of both ranks. Feature pairs were arranged in ascending order of mutual ranks, as the smaller the mutual rank, the greater the degree both features were mutually important in each other’s predictions. The formula for calculating mutual ranks is given here:

$$MR(AB) = \surd(Rank(A\to B) ✕ Rank(B\to A))$$

In the formula, MR stands for mutual rank, MR(AB) refers to the MR of features A and B, and Rank(A→B) refers to the feature importance rank which A has with respect to B. Rank(B→A) would refer to the inverse.

The top 10% of feature pairs by mutual ranks were taken to build the network. 1342 nodes and 5308 edges comprise the network. Edge weights were created by using the list of feature pairs, arranged in ascending order of mutual ranks, and inverting these ranks. These inverted ranks would then become the edge weights. Hence, if the top feature pair has a mutual rank of 1, and the bottom pair as a rank of 1 000, the top pair would be given an edge weight of 1 000 and the bottom pair would have an edge weight of 1. An inversion is done since smaller mutual ranks indicate a stronger link between features, whereas a larger edge weight indicates the same idea. As such, the constructed network is a network of features, with putative biological links between them depicted by edges.

#### Feature importance network analysis

A permutation test was used to identify statistically significantly associated feature categories, which are defined as each row of the table given in Table 2. First, the number of edges between all possible feature category pairs was calculated. Next, a permutation test with 10 000 permutations is constructed and in each permutation, features in all feature pairs are randomly rearranged, and the number of edges between all possible feature category pairs was recalculated. The number of times in which the original number of edges between categories was higher than that produced by the random shuffling of features, was recorded. This number is divided by the number of permutations to provide an empirical p-value, which identifies whether the original number of edges is significantly depleted as compared to random chance. Taking 1 minus this p-value, identifies whether the original number of edges is significantly enriched as compared to random chance. p-values were corrected for multiple testing using the Benjamini-Hochberg correction.

The feature importance network was visualised in various ways using Cytoscape 3.8.2^34^. Annotations in the network visualisation were done using a combination of manual annotations and AutoAnnotate 1.3.4^72^. Degree and betweeness centrality of nodes in the Cytoscape network were calculated by cytoHubba 0.1^73^. The overall network was depicted using the group attribute layout in Cytoscape.

#### Database development

We developed a database, finder.plant.tools (<https://finder.plant.tools>) which displays information for all features used in our study. The frontend is hosted on github and uses React.js and cytoscape.js^42^. The frontend code is available as a github repository (<https://github.com/Sweekwang/golabel>). The REST API backend uses python flask which retrieves data from Google Cloud Storage. The backend is then hosted on Google App Engine. The backend github repository is private but available upon request. The database allows the user to search for a feature, and information pertaining to it will be displayed. This information includes the general category which the feature belongs to, and a detailed description of it. In addition, feature importance scores, obtained from the trained model, will be displayed. The feature rank scores (FRS) will also be shown.

FRS is obtained by converting the feature values of the dependent variable (the feature used for predicting the target) into ranks. The feature values of the target (which is the searched for feature) will also be converted into ranks. Spearman’s correlation is calculated based on these ranks and the obtained correlation coefficient is called the FRS. This metric gives a measure of the size and direction of the association between the dependent variable with the target.

In addition to all this information, a local network is displayed. This is formed by the searched for feature, along with its first neighbours in the feature importance network. If the feature is not in the network, then the local network will not be displayed. Options to change node colour according to their feature category, filter away nodes and edges by edge weight, change network layout, and download the network in a variety of formats, are given. Edges in the local network are coloured according to their FRS values, where positive FRS values result in red edges while negative FRS values result in blue edges.
