## Supplemental Figures 1 for "Feature Importance Network reveals novel functional relationships between biological features in *Arabidopsis thaliana*"

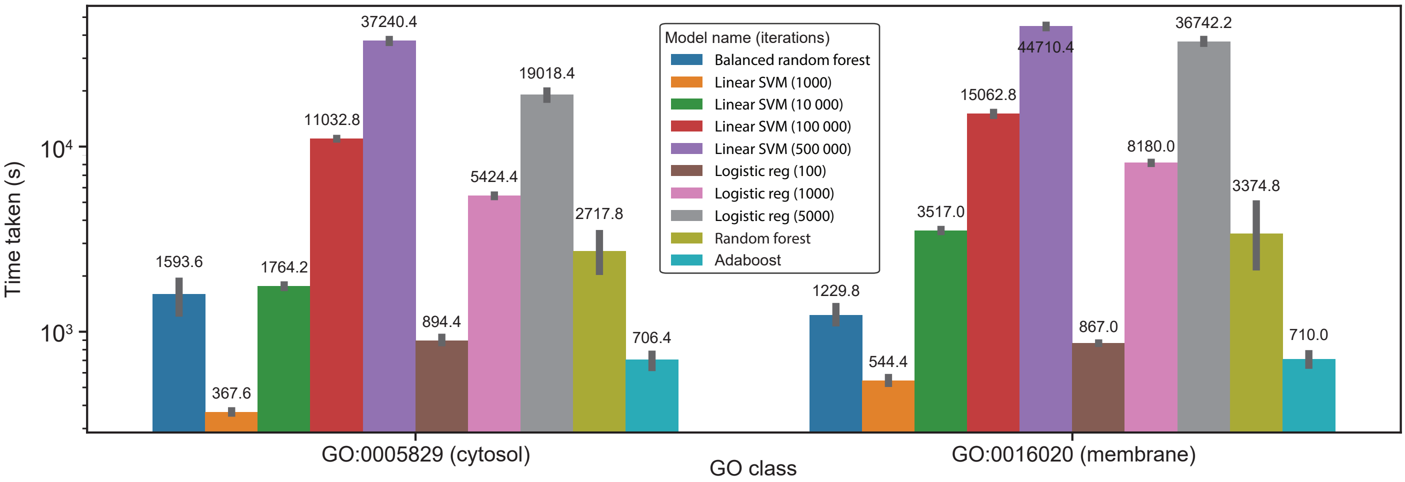


**Figure S1. Shows time taken for each individual GO class.** Time trial repeat - running all machine learning models on 2 GO classes, 5 repeats per model, to allow for all models to be compared on a common basis.

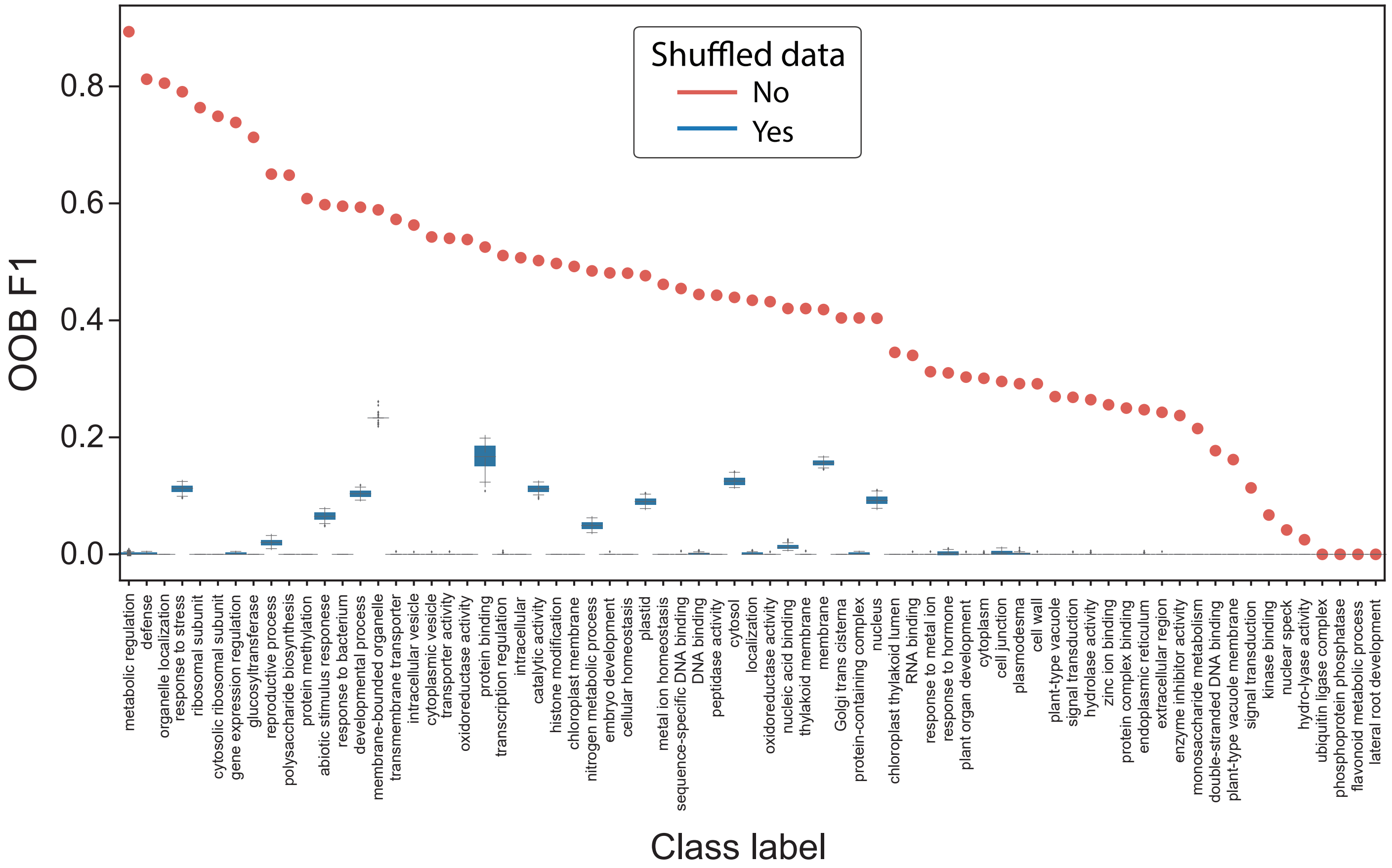


**Figure S2. Random forest OOB scores when trained on shuffled data (blue) or actual data (orange).** Some GO term descriptions have been summarised to save space, full descriptions are given in Table S9.

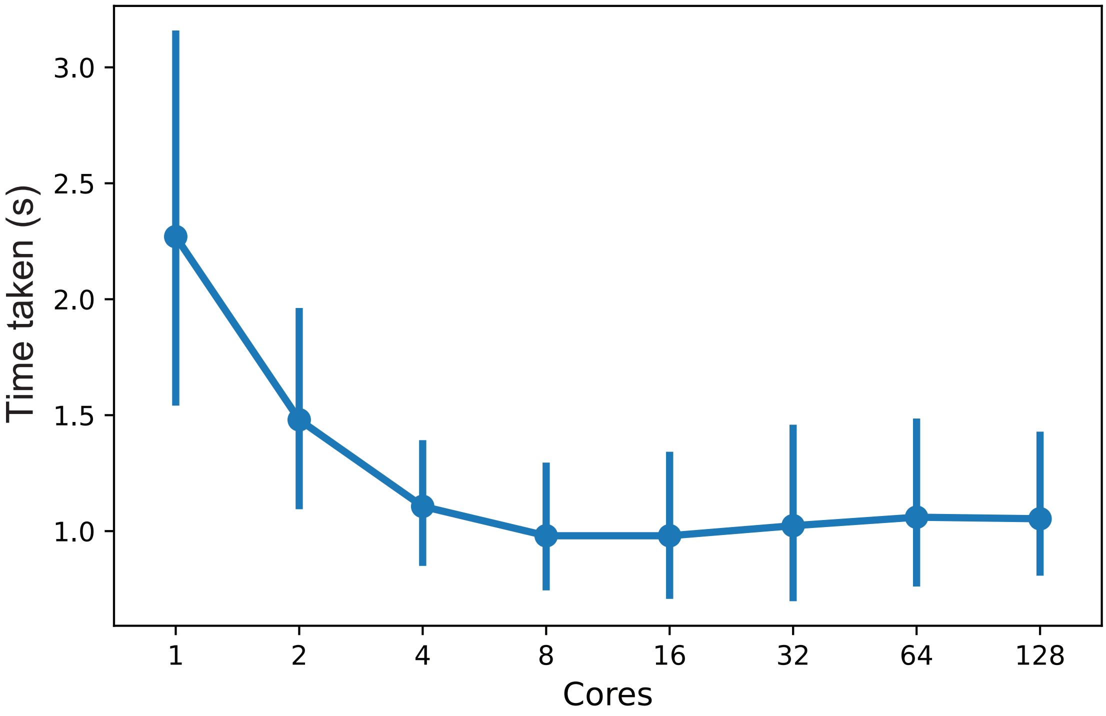


**Figure S3. Line plot showing time taken to train random forest models on 5 GO classes, GO:0043229 (intracellular organelle), GO:0005774 (vacuolar membrane), GO:0048827 (phyllome development), GO:0030001 (metal ion transport) and GO:0016836 (hydro-lyase activity).**

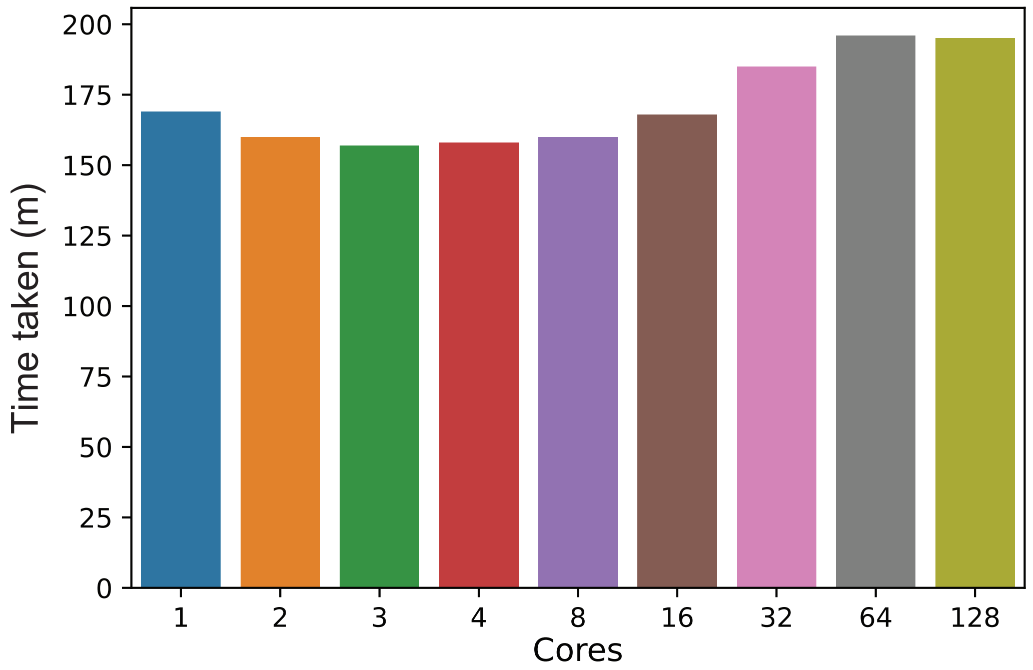


**Figure S4. Bar graph showing time taken to train random forest models on all GO classes.**
